# Deliberation and enaction during adaptive economic choice

**DOI:** 10.1101/445346

**Authors:** David J. Hawellek, Kevin A. Brown, Bijan Pesaran

## Abstract

Economic decisions can adapt to contexts. Choices can be quick and impulsive or slow and more deliberative, depending on the temporal context. Choices can also depend on how we enact the choice, in an action context. Where we decide to go for dinner may change if we can take a taxi or need to walk.

We hypothesized that frontal action circuits could contribute to adapting economic choices to context because of their privileged position over actions as endpoints of decisions.

To test this, we performed an unbiased survey of neuronal population activity across motor, premotor and prefrontal cortices as animals expressed context-dependent economic preferences. Activity in distributed action circuits tracked the animals’ evolving preferences in real-time and integrated them with a signal for their enaction. We propose that frontal action circuits form a neural substrate that supports an adaptive control over economic choice by flexibly translating real-time preferences into actions.

## Introduction

When we make economic decisions, we express subjective preferences. Forming preferences requires a deliberation process that compares different options. Expressing a preference requires an enaction process that translates the deliberation into choices. Together, these two processes – deliberation and enaction – control our decisions. A central feature of our goal-directed behavior is that the control over our decisions is flexible and allows decision making to adapt to contexts^1,2^.

Different temporal contexts often profoundly affect our decisions. Faster decisions tend to be objectively less optimal than slower decisons. This is often observed as a progression from impulsive to more deliberative and even optimal choices as time goes by^3–7^. How we enact decisions also provides an important action context. Our preferences can change depending on how we express them. For example, decisions made with a reach require effort and can show strong spatial biases^8^, while saccades are comparatively effortless and reveal different decision biases^9–11^. In this way, how we enact a choice can itself become part of the decision process. A key question is how neural circuits allow decisions to adapt to changing temporal and action contexts.

Action circuits extend across the frontal and prefrontal cortices and are well situated to track action preferences and translate them into choices in real-time^12–14^. We hypothesized that action circuits could in this way contribute to adapting choices to contexts. Specifically, if a neural circuit supports context-dependent choices, then activity that reflects a preference in one context should reorganize to reflect a different preference as the context changes – a signature of deliberation. Furthermore, to enable flexibility over time, the neural activity should reflect the resolution about whether it is the right time to translate a preference into a choice – a signature of enaction.

We trained two monkeys to perform a dynamic decision task in which we manipulated the temporal and action contexts of decisions. Temporal context was manipulated by either instructing animals to make their choice immediately or to wait and make their choice after a delay. Action context was manipulated by either instructing animals to make their choice with a reach or a saccade. Behaviorally, each animals choice depended strongly on context. Reach choices were qualitatively different when made immediately or after a delay. Biased and suboptimal immediate reach choices became unbiased and more optimal delayed choices, as if deliberation led to more optimal decisions. In contrast, saccade choices depended less on the temporal context and were more optimal overall. Thus, depending on the action, the preferences of the animals evolved dynamically, akin to an unfinished and continuous deliberation that could be quickly and flexibly translated into choices at different points in time.

We performed a large-scale survey of neuronal ensemble activity by systematically making distributed recordings across the dorsal surface of frontal and prefrontal cortices. In line with our hypothesis, distributed frontal ensembles exhibited signals related to the deliberation and enaction that supported the context-dependent choices of the animals. Action related circuits, prominently the dorsal premotor cortex, tracked the preferences of the animals in real-time and combined them with an enaction signal that could flexibly propagate preferences into choices. In sum, we contribute key evidence to support the idea that frontal action circuits can exert control over economic choice that allows adapting choices to contexts.

## Results

### Dynamic decisions with common real-world constraints

We trained two monkeys to perform a **dynamic foraging task** with three critical components (**Fig. 1a**). First, we unpredictably forced the animals to make a decision immediately rather than after a short delay on a third of the trials. Thus, while there was usually a brief moment to come up with a choice, the animals sometimes had to immediately commit to an option. In this way, we introduced two different temporal contexts for making decisions. Second, the reward magnitudes at the movement targets continuously drifted in an uncorrelated way. Thus, the decision behavior relied on continuous learning and could not be based on simple deterministic strategies. Third, the animals sampled the identical environment with either saccades or reaches. The oculomotor system operates under different constraints than the skeletomotor systems. Specifically, saccades are fast and energetically less costly. In this way, we studied how decisions could depend on two different action contexts.

**Figure 1:**
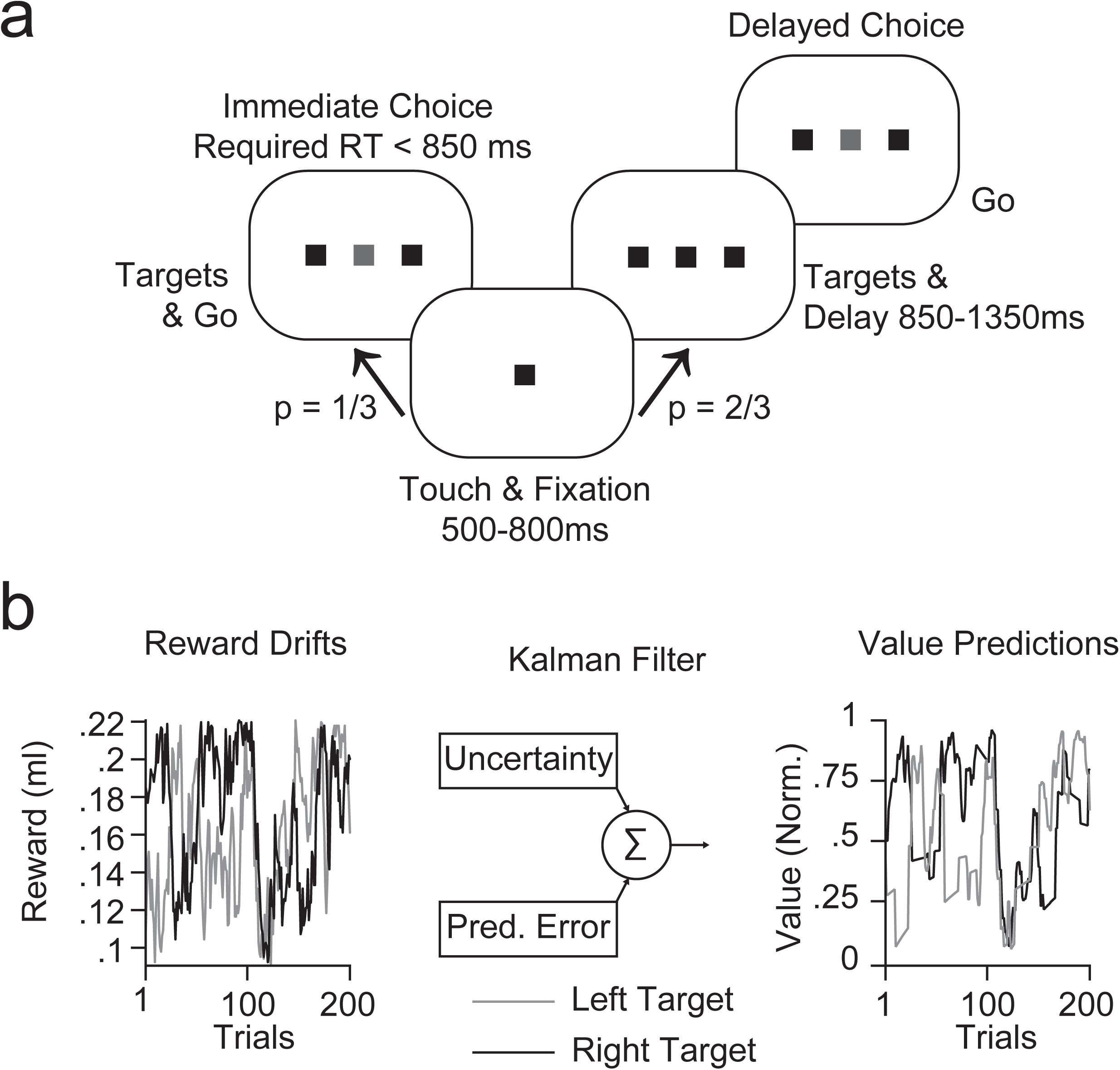
Dynamic foreaging task. a) Animals started a trial by acquiring central touch and fixation. Two diametrically opposed movement targets appeared on a horizontal axis. On 2/3 of all trials the animals had to wait until the dimming of the central stimulus indicated signal to go (Delayed choice). The animals were free to choose either movement target to obtain a fluid reward. On 1/3 of all trials the go cue was given immediately with the appearance of the choice targets. On immediate choice trials the animals had to achieve a reaction time that was shorter than the shortest delay on delayed choice trials. Animals made choices either with a reaching movement while maintaining fixation or with a saccade while maintaining touch. In all subsequent analyses the term ipsilateral is used to refer to the side of the reaching arm. The reach and saccade conditions alternated in longer blocks. b) The fluid magnitudes at the two targets drifted across trials in an uncorrelated way. We matched the statistics of the drifts across days and actions by replaying drifts after a few days, exchanging the association of target and drift and counterbalancing the sequence of reach and saccade blocks (**see Methods**). We used a Kalman filter to predict value states of the targets based on the animals’ choice behavior. These values are referred to as objectively optimal values as detailed in the main text.

Our goal was to study how neural computations allow economic choice to adapt to such varying contexts, achieving a flexible control of the decisions.

### Animals dynamically compute context-dependent preferences

A computational model of choice behavior revealed that animals made qualitatively different reach decisions according to the temporal context. We first used a reinforcement learning framework to infer values from rewards offered during the two-arm bandit decision task^15^. The model obtained predictions about the optimal trial-by-trial value states of the movement targets by using a Kalman filter to predict the sequence of choices from experienced rewards (**Fig. 1b**). The value predictions made by the computational model correspond to those of an ideal observer and are optimal in that they represent an upper bound on what the animals could have learned through sampling. We refer to these predictions as the objectively optimal values.

With this definition of optimality in hand, we observed that animals made qualitatively different decisions depending on both the action context and the temporal context. We found that delayed reaches (DR) were more optimal than immediate reaches (IR). IR were less successful than DR (Hit rate. Monkey J: IR 63.5%, DR 67.1%, p = .02; Monkey H: IR 69.4%, DR 73.6% high value choices, p = 0.004, Wilcoxon rank-sum tests). The more optimal behavior of the DR was associated with a change in the reach bias of the animals across temporal contexts (**Fig. 2a, S1**). While IR were biased and suboptimal, choices became gradually more evenly distributed across the two movement options, achieving an objectively more optimal income for DR. Since the temporal context was unpredictable trial-by-trial, this suggests animals revised their choices in real-time, consistent with a form of ongoing deliberation.

**Figure 2:**
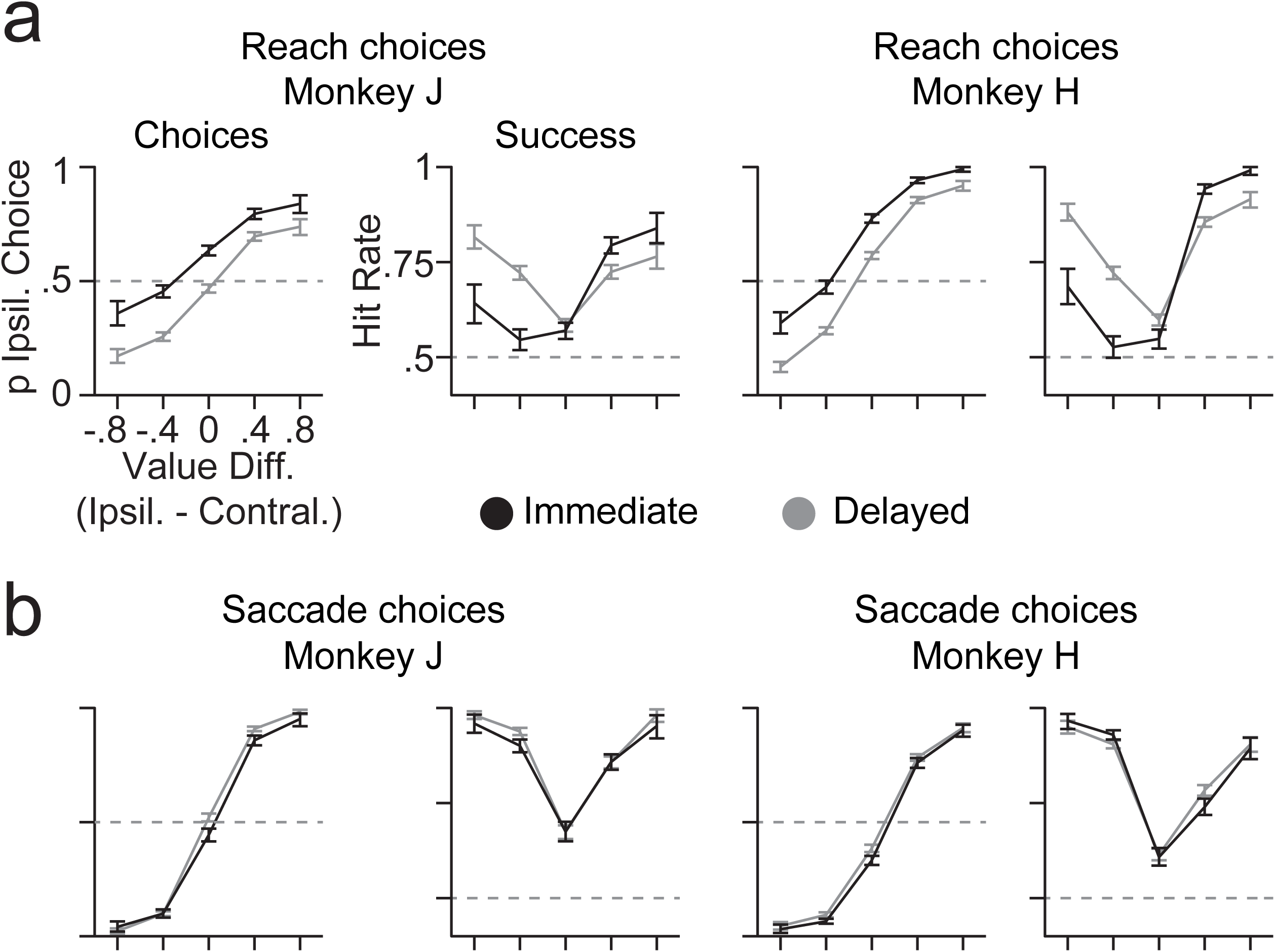
Animals change their reach preferences dynamically. a) Choices and success for reach decisions of monkey J and monkey H. IR (black) and DR (grey) choice probabilities are shown according to the objectively optimal values at the two target locations. Positive value differences indicate a higher value at the target ipsilateral to the reaching arm. Success is depicted with the hit rate that reflects the probability of choosing the objectively higher valued target for the same choices. Dotted lines depict the probability of choosing both sides equally as well as the chance level of detecting the objectively optimal reward. All error bars depict 95% confidence intervals. b) Same as a but for saccade choices.

In comparison, saccade decisions were objectively more optimal than reach decisions (Overall percentage of hit trials, Monkey J: Reaches 65.3%, Saccades 82.5%, p < 10^−8^; Monkey H: Reaches 71.3%, Saccades 77.3%, p = 1.4×10^−8^, Wilcoxon rank-sum tests). Saccade decisions also exhibited a more stable pattern of choices in the two different temporal contexts of immediate and delayed saccades (IS, DS, **Fig. 2b, Fig. S2**).

More detailed logistic regression models of the choice behavior based on the Kalman-inferred values and behavioral timing parameters such as reaction times corroborated the evidence for context-dependent decision making. (**Fig. S1, S2, Supplementary Results**).

Overall the choices of the animals were strongly influenced by the temporal and action contexts of the decisions. Most strikingly, reach choices and not saccade choices were associated with a dynamic change in choice preferences of the animals over time, with qualitatively different decisions made immediately or after a brief delay. These behavioral observations are consistent with the idea that the context-dependent choices are revised online. This indicates that making choices involved a process of ongoing deliberation that was dynamically read out through enaction.

We next measured neural dynamics in frontal and prefrontal cortices to test how neural activity that predicts choice in one context reorganizes as the context changes.

### Multiplexed coding of choice and enaction by distributed cortical ensembles

We simultaneously recorded spiking activity across a 2 cm extent spanning the frontal and prefrontal cortices using a semi-chronic multielectrode array (SC96, Gray Matter Research, USA, Monkey J: 354 single units, 753 multi units; Monkey H: 96 single units, 664 multiunits). The array covered pre- and post-arcuate cortex including prefrontal cortex (PFC), the dorsal and ventral aspects of premotor cortex (PMd, PMv) as well as parts of somato-motor cortex (SM, **Fig. 3a,** N units: Monkey J, SM=143, PMd=568, PMv=100, PFC=296; Monkey H, SM=81, PMd=227, PMv=132, PFC=320). Consistent with known anatomical connections and functional specializations, neural activity across the frontal cortex (SM, PMd, PMv) responded strongly during the immediate and delayed reach tasks (**Fig S3**) while activity across the PFC most strongly responded during the immediate and delayed saccade tasks (**Fig S4**).

**Figure 3:**
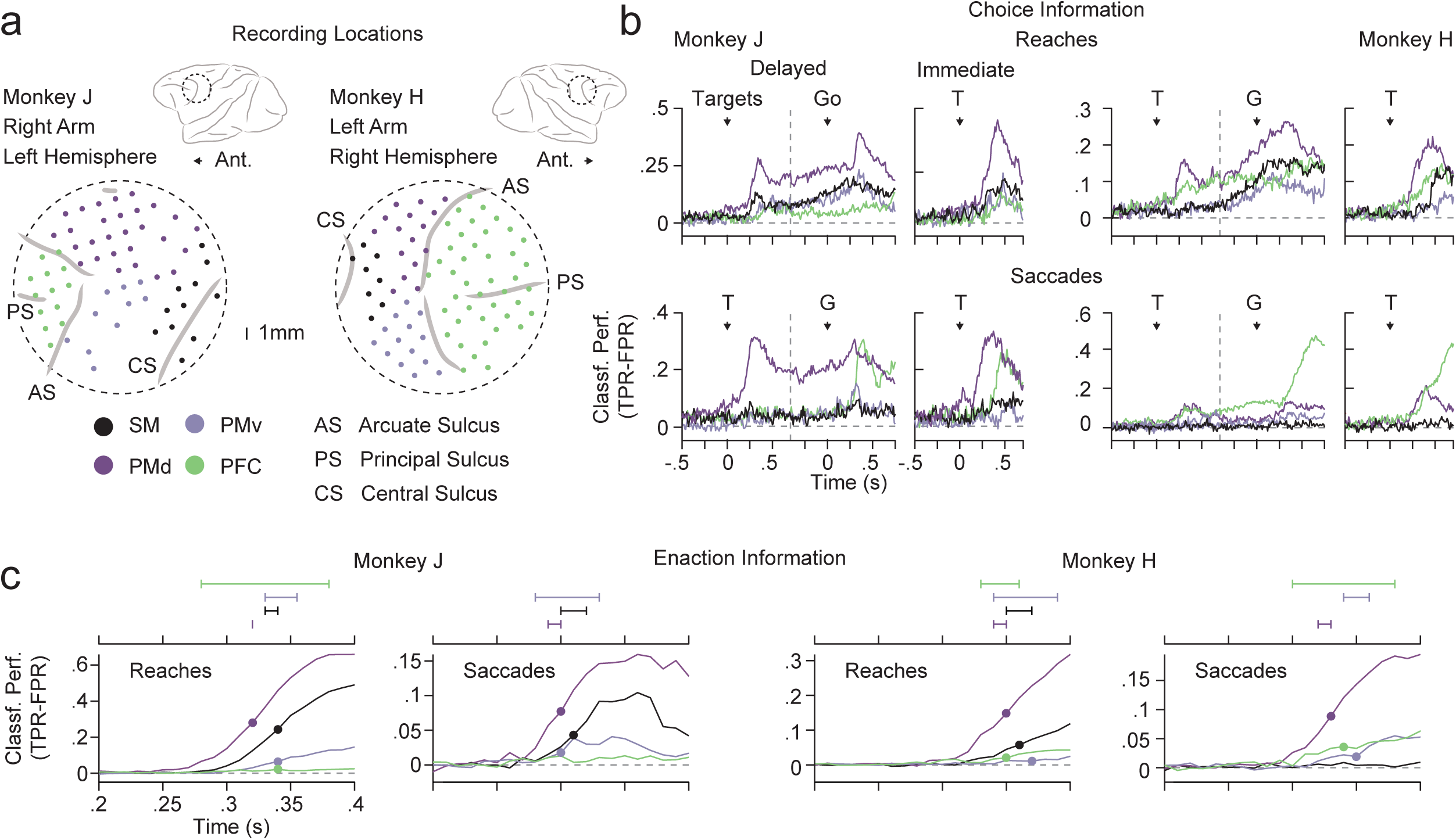
Frontal ensembles combine information about choices and their enaction. a) Recording locations. The drives were centered on the spur of the arcuate sulcus covering pre- and post-arcuate sulcus. Recordings were made in the hemisphere contralateral to the reaching arm. Anatomical locations were derived from co-registration of MRI data to the implanted drives. Grey lines depict sulci. PS, principal sulcus, AS, arcuate sulcus; CS, central; sulcus. Electrode locations are shown as colored dots. Only electrodes that contributed spiking activity to the analyses are shown. Electrode colors depict anatomical subdivisions. SM, somato-motor; PMd, dorsal premotor cortex; PMv, ventral premotor cortex; PFC, prefrontal cortex. T, target onset; G, go cue. b) Classification performance of ipsilateral and contralateral choices based on spiking activity within the anatomical subdivisions (choice information). TPR, true positive rate; FPR, false positive rate. The left column of panels depicts data for monkey J, the right column depicts data for monkey H. Classification performance is shown separately for delayed and immediate choices as well as reaches and saccades. The data for delayed choices is shown for two alignments, to the target onset as well as the go signal. The transition between the two periods is indicated as a vertical dotted gray line. Note that all analyses were based on the spiking activity of the same neurons. c) Classification performance of immediate and delayed trials based on spiking activity within the anatomical subdivisions (enaction information). Only data up to 50 ms before movement onset was used to perform classification of immediate and delayed choices. The time axis is chosen to show a more detailed view onto the time after target onset. We estimated latencies of information onset as the time point at which the information reached half maximum within the shown time window. Latencies were only estimated if significant classification performance was detected. Small panels above the line plots depict the 95% confidence interval of the latencies for each anatomical subdivision obtained by bootstrapping. The median latency is depicted as a circle on the classification performances for the anatomical subdivisions.

We used multivariate classification (Linear discriminant analysis - LDA) based on spiking activity to track information about upcoming choices as well as the pending enaction of a choice. We decoded trials based on the animals’ upcoming movement choice (choice information, **Fig. 3b**) or the temporal context, i.e. whether a choice had to be enacted immediately (enaction information, **Fig. 3c**). Both types of decision related information were widely distributed and could be decoded from nearly all anatomical subdivisions before overt behavior (**Fig. 3b, c, Fig S3, S4**). We generally observed that the classification performance for saccades compared to reaches was lower (**Fig. 3b,c**), except for PFC which showed more pronounced activity and more information for saccades (**Fig. S4**).

A prominent feature during both reach and saccade choices was a brisk global increase in spiking activity after the onset of the choice targets (**Fig. S3, S4**). This increase in activity was evident across all cortical areas independent of spatial choice or temporal context. The choice information exhibited similar early increases, even for delayed choices for which the enaction of a choice was withheld for the delay. This pattern was particularly pronounced for PMd and was present for both action contexts (**Fig. 3b**). While choice information increased, neural activity also started to diverge depending on whether or not a decision had to be enacted immediately (**Fig. 3c**). Neural activity in PMd again robustly contained such enaction information, leading a cascade across areas (**Fig. 3c, Fig. S3, S4**). Notably, PFC contained only little enaction information, again, despite the slightly varying coverage between the two animals (**Fig. 3a**) and the reach or saccade context of the decisions (**Fig. 3c**).

In sum, we observed distributed, dynamic and action context dependent representation of decision related information. Neuronal ensembles tracked information about upcoming choices along with an enaction signal supporting the flexible allocation of choices.

We next considered how choices were revised in real-time by comparing immediate and delayed choices. What turns suboptimal IR into more optimal DR?

In the dynamic foraging task, rewards for each option change on each trial. The animal’s choices may have depended on how long it takes to compute the objectively optimal value for each option. Information about the objectively optimal values may have been weak at the beginning of a trial, built up slowly over time and have been strongest around the time of delayed choices, explaining the revision of choices across temporal contexts. We next investigated the influence of the objectively optimal values on the ensemble dynamics.

### Objectively optimal values influence ensemble dynamics early

We pooled simultaneously recorded units into array-wide ensembles to capture the distributed nature of decision related information (**Fig S5a, b, Supplementary Results,** Monkey J: 29 ensembles, median size 22 units, Monkey H: 27 ensembles, median size 23 units).

We predicted that a larger difference in the Kalman-inferred values between the two targets would lead to better choice classification performance. Specifically, if the objective values were driving the revision of the animals’ choices over time, the correlation between single trial value differences and classifier predictions should increase over the course of delayed choice trials. To test this prediction, we regressed the single-trial value differences against single-trial classifier probabilities obtained in 500ms windows centered at four intervals of interest during delayed choice trials (**Fig 4**; Baseline, Commit, Pre-Go, Reaction Time). We defined the commit time point as the local peak in classification performance within the first 500 ms during delayed choices, i.e. when choice information started to separate depending on whether or not animals enacted a choice. To avoid the confounding influence of choice signals, we performed the regressions separately for ipsilateral and contralateral choices and then averaged the results (**see Methods**).

**Figure 4:**
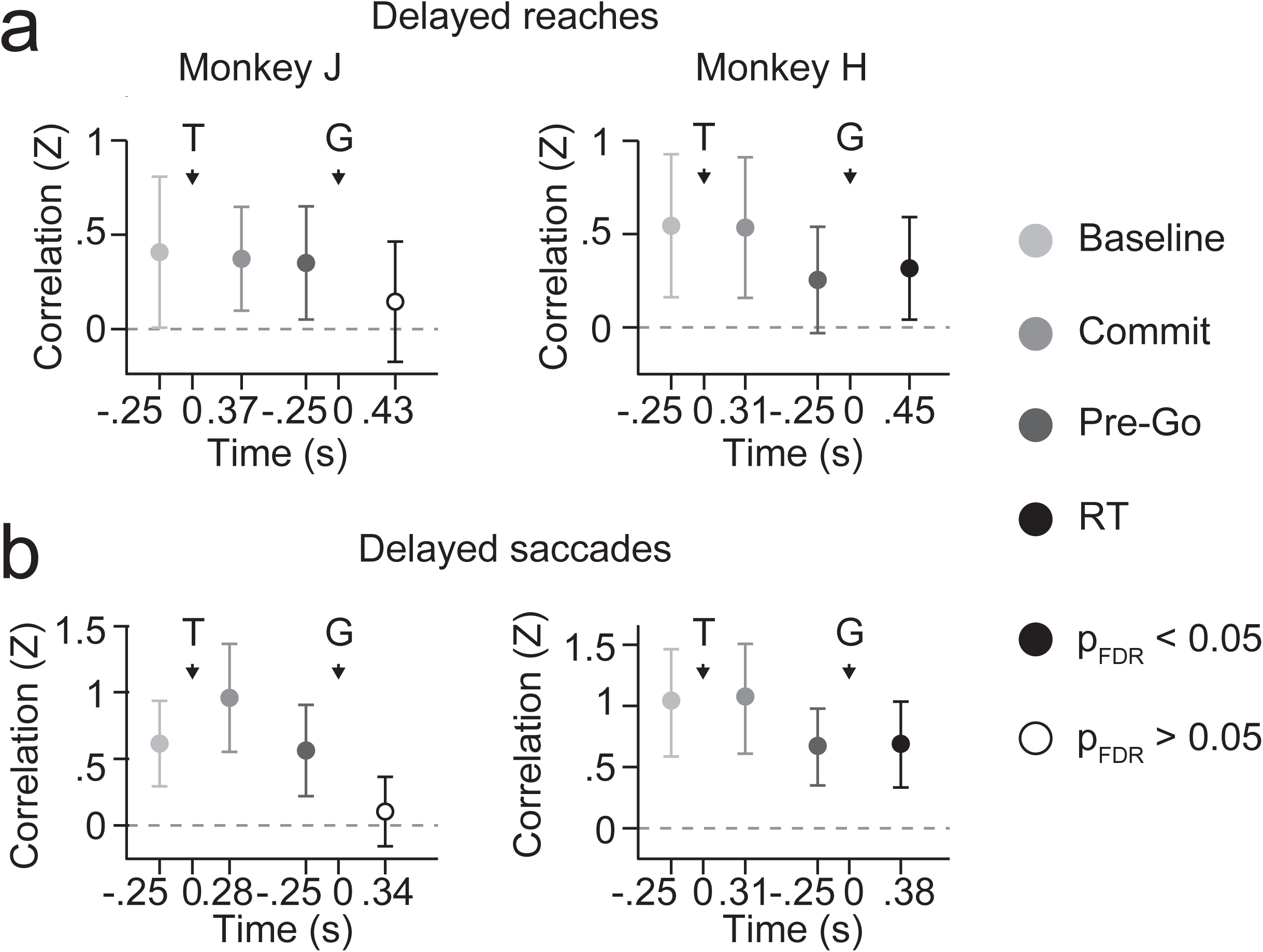
Objective values influence ensemble dynamics early. a) Relationship of single trial value differences and classifier probabilities at four time points during delayed reaches (DR). Correlations are shown as z-scores relative to a resample distribution obtained through random permutations (**see Methods**). Significant influences of the objective values on the classification performance are depicted as solid circles. All error bars depict 95% confidence intervals. The baseline window was chosen as the 500 ms preceeding target onset. The commit window was chosen as 500 ms centered on the time point at which choice information peaked during delayed trials before then slightly decreasing. At this time point information continued to rise on immediate choices, signaling the pending enaction. The pre-go window was chosen as the 500 ms preceding the go cue. The reaction time (RT) window was chosen as 500 ms centered on the median reaction time. T, target onset; G, Go cue. Note that all correlations were performed separately for ipsilateral and contralateral choices to avoid choice signals confounding the analysis. b) Same as a for delayed saccade (DS) choices.

The Kalman-inferred values significantly influenced the classification of choices throughout the delayed trials for both reaches and saccades (**Fig 4**, t-tests, p < 0.05, FDR corrected). Strikingly, the regressions were significant already during the baseline intervals and remained at comparable levels up to the movement period. This observation is in line with the idea that animals predicted the value states of targets in real-time and encoded the optimal value states of the targets at the start of the trial. Further in line with this observation is the finding that saccade choices exhibited a more stable influence of the objective values throughout the trial (**Fig 2, Fig S2**). Thus, the process of learning and updating objectively optimal values was fast enough to guide the even faster saccade behavior early on and in a stable way. These results do not support the view that immediate reach choices were suboptimal because information about the Kalman-inferred values built up only slowly in the course of a trial.

Next to changes in objective value signals, choice revisions may result from a dynamic change in other bias signals that together constitute the animals’ subjective preference. Such bias signals may act on pools of neurons that represent the choice options simultaneously and compete over the mutually exclusive selection for choice^16^. We next investigated whether subjective preferences were tracked through two circuits that represented the choice options.

### Action circuits dynamically track subjective preferences

We split the neurons into two groups that encoded the two alternative movement directions and assessed the evolution of their activity during the different choices (**Fig 5, see Fig S6 and Fig S7**).

**Figure 5:**
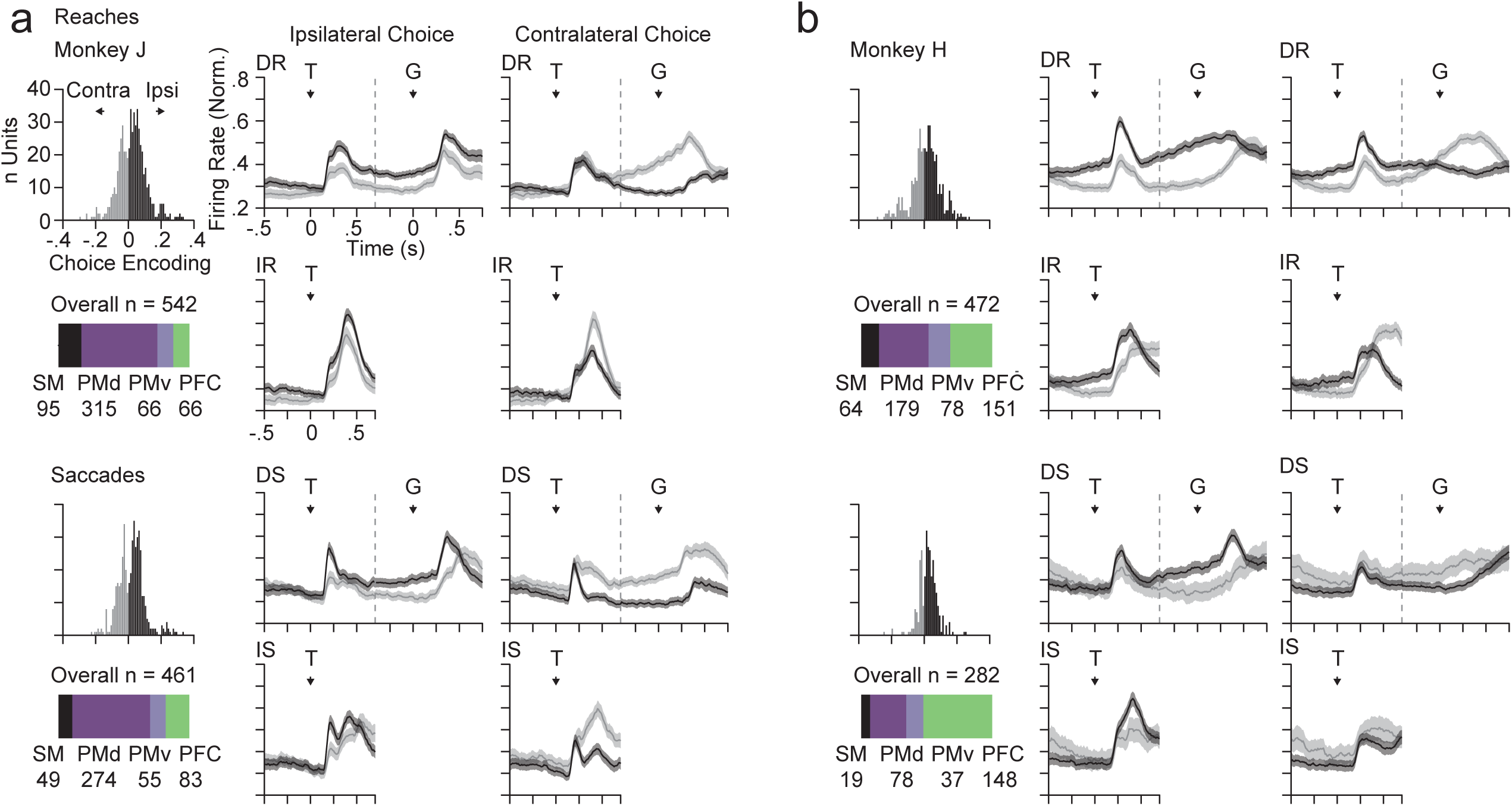
Action preferences are tracked through competing circuits. a) Activity profiles of two pools of neurons encoding ipsilateral and contralateral choices through increases in their firing rate. The histograms show the encoding strength of the two pools of units computed as the AUROC minus 0.5 with ipsilateral choices as consistent positive class. Each unit with negative values increases its firing rate for contralateral choices, each unit with positive values increases its firing rate for ipsilateral choices. Units were selected for the analysis for having a significant difference in their firing rate between ipsilateral and contralateral choices in the 500 ms preceding delayed choice reaction times (p < 0.05, ranksum test). Units were then split into the two pools based on the sign of their choice encoding. The colored bar depicts how many neurons of each anatomical subdivision contributed to the analysis. The line plots show the activity of each pool of neurons for ipsilateral (left column) and contralateral (right column) choices as well as immediate (top row) and delayed (bottom row) trials. Spiking activity was normalized by dividing with the maximum activity across all conditions for each neuron separately. DR, delayed reaches; IR, immediate reaches; DS, delayed saccades; IS, immediate saccades. T, target onset; G, go cue. All shadings depict 95% confidence intervals. b) Same as A for monkey H.

In line with a competition over selection for enaction, the two pools of units exhibited dynamic activity profiles of that reflected the animals’ choice allocation and tracked their subjective preferences. For reaches the pool of units encoding ipsilateral movements was enhanced early during the trials during the baseline and up to the commit, agreeing with the ipsilateral behavioral bias for immediate reaches (**Fig 6**). For delayed reach choices, the two pools of neurons gradually evolved toward a more balanced state reflecting the more balanced allocation of choices across the two choice options.

**Figure 6:**
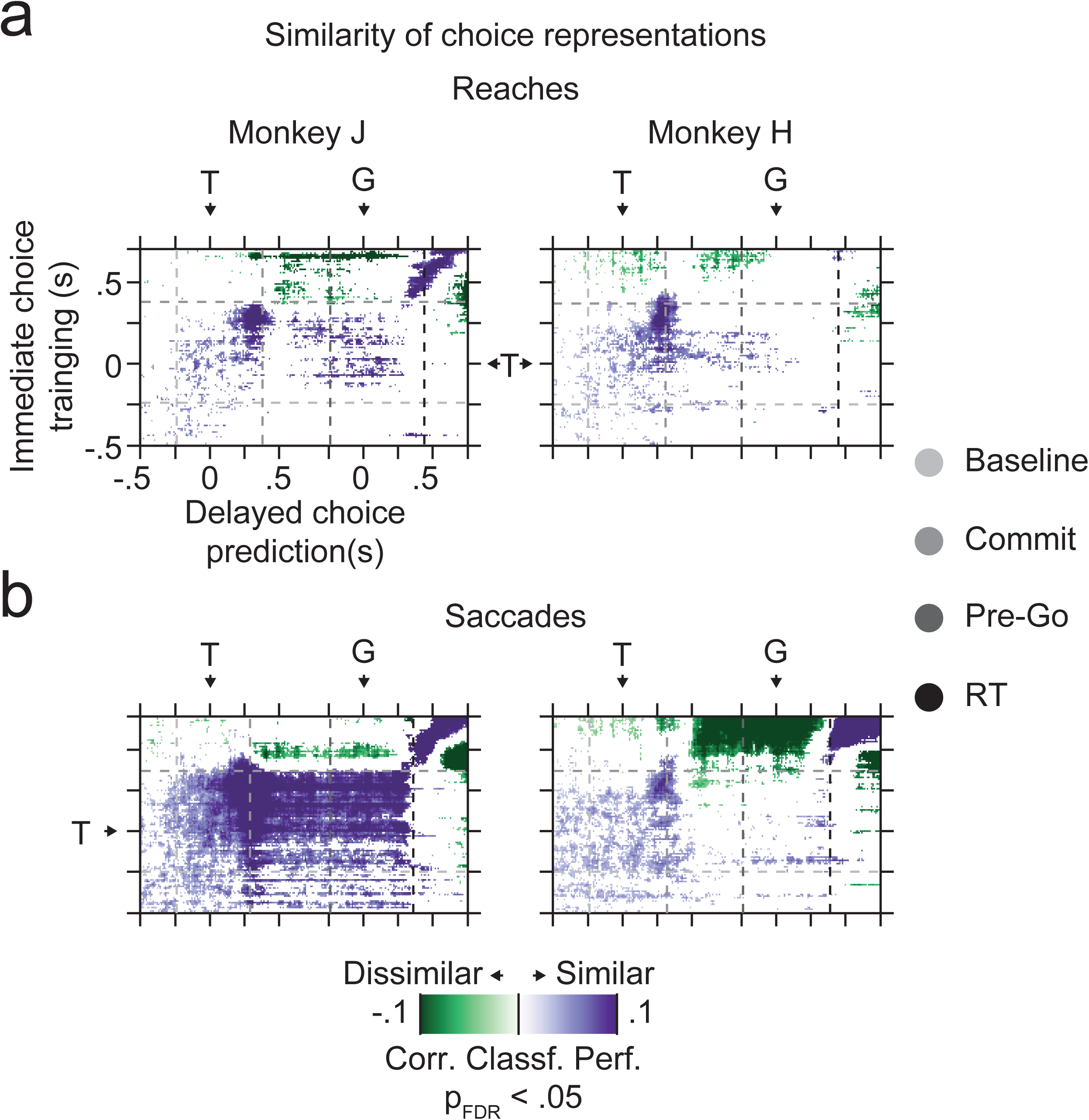
Similarity of the representation of choice across array wide ensembles. a) We trained choice classifiers on immediate reach trials and predicted delayed reach choices based on activity in delayed reach trials for different combination of time points. The resulting cross-classification was corrected by subtracting the pattern of classifier performance that would be expected based on the product of training and predicting within immediate or delayed trials, respectively. Negative values indicate a consistently weaker and positive values a consistently better than expected cross-classification performance, indicating dissimilar (green) or similar (purple) neuronal representations between immediate and delayed choices. Dotted lines indicate the four time points of interest throughout the trials for orientation. T, target onset; G, go cue. The data is masked to only show significant time points as colored (p < 0.05, FDR corrected).

For monkey J’s saccade decisions, the activity profiles were more balanced from the beginning of the trials, matching the choice behavior. The brisk increase of activity around the commit was still slightly more pronounced for the ipsilateral pool of neurons, agreeing with a slight ipsilateral choice bias for very fast immediate saccades (**see Fig S2**). Similarly, early enhanced activity of the pool encoding contralateral movements for monkey H’s saccades matched a contralateral choice bias.

Overall, context-dependent preferences of the animals were tracked through the activity of competing circuits that represented the available choice options. In other words, the process of deliberation could be read out from action circuits in real-time.

Fast and suboptimal and slow and more optimal decision making is a well described phenomenon in behavioral economics and psychology^3–5,7,17^. An idea permeating many models of such decision behavior is that dynamic changes in decision making are supported by a transition between different decision systems. In principle, the choice revisions here may have resulted from a transition between two decision systems over time. For example, between a suboptimal model-free system that is biased and/or depends on heuristics and another computationally more demanding model-based system. A prediction of such dual decision system models is that the representational structure of choice across the neuronal ensembles should change over time. We next investigated whether a change in the representational structure across the neuronal ensembles may have been associated with the choice revisions from immediate to delayed decisions with a cross-classification approach.

### Context-dependent choices are not associated with a transition between decision systems

We trained classifiers on the ensemble activity at different times during immediate choice trials and predicted delayed choices based on ensemble activity at different times during delayed choice trials. We hyothesized that a transition between decision systems should lead to a difference in the ability to predict delayed choices based on immediate choices.

We found significant cross-classification performance between immediate and delayed choices that was changing over time (**Fig S9,** p<0.05, permutation tests, FDR corrected). Cross-classification ramped up during the baseline period to peak at the commit time point and reaction time periods. In order to assess whether such changes in cross-classification reflect a change in the underlying choice representations, the dynamic changes in classification performance within the immediate and delayed trials need to be accounted for (signal-to-noise ratio). The pattern of cross-classification expected to result purely from the changes in the individual classifier performances is predicted by the product of the classifier performances within immediate and delayed choice. We computed a corrected cross-classification performance (**Fig 6**) by subtracting this expected pattern from the raw cross-classification performance of each ensemble. The corrected cross-classification revealed both similarities and dissimilarities in the choice representations of immediate and delayed choices (**Fig 6**, p < 0.05, FDR corrected, permutation tests). A consistently positive corrected cross-classification indicates a surprisingly strong cross-classification performance and suggests a similarity of the representations between immediate and delayed choices. A consistently negative performance indicates a surprisingly weak cross-classification, suggesting a dissimilarity of the underlying representations.

A central pattern visible for reaches as well as saccades was that the commit time point reflected a moment in time at which the representational structure of choice pivoted from an early similar representation, present just after the baseline, to a later dissimilar one towards choice execution (**Fig 6a, b**). This change in representation is in line with the idea of transition from deliberation to enaction. Furthermore, after a choice has been withheld at the commit time point during delayed trials, the representational structure regained similarity with earlier activity. These observations suggest that the neural activity reverted to earlier states during the delay. Importantly, we did not observe significant dissimilar representation when comparing the time after the commit for DR and before the commit for IR. That is, there was no evidence for a change in the representational structure between IR and DR, providing evidence against a transition between decision systems for the revised choices. Instead, IR and DR choices were represented similarly across the recorded ensembles. We repeated the analyses for predicting immediate choices when training on delayed choices and found highly similar patterns (**Fig S9**)

An observation of note is that monkey J’s saccade choices that were associated with the most stable and most strongly objective value driven decision behavior (**Fig S2, Table S1**) were associated with the most widespread and robust pattern of cross-classification similarity between immediate and delayed choices (**Fig 6b**). Such stable and robust cross-classification may thus be a signature of a highly stable deliberation process. Along the same lines, the reach cross-classification of both monkeys as well as the saccade cross-classification of monkey H was less pronounced. Accordingly, the behavior of all these were best modelled containing inversions of reaction time dependent biases (**Fig S1, S2, Table S1**), suggesting the presence of more complex dynamics with sensorimotor processes contributing to the decisions.

## Discussion

Contexts play a central role when primates make decisions. A key question is how neural circuits allow adapting choices to contexts to enable flexible behavior.

We find that action circuits multiplex decision signals that reflect a subjective deliberation over actions in real-time and the selection of a choice through enaction^18–20^. When reaching, animals changed their choice preferences on brief time scales such that suboptimal immediate choices turned into more optimal delayed choices. These observations are consistent with the idea that a continued deliberation may have flexibly been read out through the enaction of choices if necessary. Such fast and flexible gating of current action preferences may constitute a form of context-dependent control over economic choice at a stage that connects deliberation with enaction. In other words, we observed signatures of control over economic choice most proximal to the endpoint of decisions – action.

Previous research in more static decision environments highlights the role of higher level cortical regions such as e.g. the orbitofrontal cortex in the control over economic choice^11,21^. An observation consistent with the idea that high level valuation circuits may fully determine economic decision processes were the saccade choices of monkey J. Monkey J’s saccade choices were highly stable over time and reflected the most faithful enaction of objectively optimal values in our data^1^. Such a stable decision process may be most strongly driven through the valuation circuitry. An important implication of the reach decisions reported here is that action contexts may be instrumental to reveal more complex and action dependent choice mechanisms in which actions are part of the decision problem. The computations that support choice are distributed^2,12,22–25^ and extend across several circuits that could provide complementary control over choices in a context-dependent way. Our results are in line with the idea that the context of decisions may define the exact way in which neural circuits exert their control over decisions^2^.

The large-scale electrode arrays allowed for a centimeter scale coverage of peri-arcuate cortex. Nonetheless, a central limitation to our study still the scope of the neural recordings. The contribution of other prefrontal, parietal or subcortical brain structures such as the basal ganglia to the dynamics described here remain unclear, yet. The basal ganglia may supply temporal signals during dynamic decision making that likely constitute a central component of the process of enacting choices^19^. Thus, the full neural mechanism that mediates the flexible gating of choices as observed here may involve more extensive subcortical and cortical networks outside of the scope of our arrays.

Along the same lines, we found that the suboptimal immediate and more optimal delayed reach choices were not associated with a change in the representational structure across the neural ensembles. Such a change in choice substrate for the qualitatively different decisions would have been in line with theories that assume multiple decision systems to underlie the dynamic control of behavior^3–5,7,17,26^. Our results instead suggest that the frontal ensembles contained a more unitary substrate to support the different choices, similar to findings that show a surprising overlap in striatal substrates for behaviorally dissociable types of decision making^27^. However, our findings do not exclude the possibility that the frontal ensembles here constitute a point of convergence of separate decision systems for the immediate and delayed reach decisions that extend across other parts of the brain.

The observation that the objective values influenced the ensemble dynamics similarly throughout the trials suggests that factors that are intrinsic to reaching were dynamically adapted over the course of delayed trials to drive differences in decision making. Although the exact nature of these intrinsic bias signals remains unclear, one parsimonious explanation is that the animals dynamically discounted action costs during delayed reach trials to countermand the ipsilateral decision bias. The activity across the frontal ensembles tracked subjective preferences in a way that aggregated such intrinsic changes together with the extrinsic (objectively optimal value) decision factors. An important question for future investigation is how dynamic decision signals intrinsic to actions arise. Motor cortices may present a suitable substrate for the real-time construction of such dynamic action variables themselves. Alternatively they may form loops with other prefrontal cortical regions such as the orbitofrontal cortex or regions that e.g. exhibit effort related signals such as the anterior cingulate cortex^28^ to dynamically control subjective action variables. Importantly, the idea that the frontal ensembles tracked multifaceted preferences in real-time suggests that these circuits are a formidable substrate to support choices that quickly adapt to different decision contexts.

In sum, temporal and action contexts strongly impacted the way that non-human primates made decisions. Action circuits exhibited signatures of decision control by combining signals related to deliberation and enaction enabling a flexible adaption of choices to different contexts.

## Methods

### Experimental preparation

Two male adult rhesus macaques *(Macaca mulatta)* participated in the study. Monkey J weighed 10 kg and Monkey H weighed 12 kg. All surgical and animal care procedures were approved by the New York University Animal Care and Use Committee and were performed in accordance with US National Institute of Health guidelines for care and use of laboratory animals. The animals had participated in electrophysiological experiments targeting the parietal cortex previous to the experiments presented here and have been trained consistently with established training protocols for this study. Before behavioral training we implanted head-restraint prosthesis. Each monkey was trained in an unlit sound-attenuated electromagnetically shielded room (ETS-Lindgren). Following behavioral training we implanted customized recording chambers (Gray Matter Research, USA) over the spur of the arcuate sulcus using image-guided sterotaxic surgical procedures (BrainSight, Rogue Research, Canada). The chambers and microdrives were customized to accommodate the individual anatomy of the animals. During surgery, we measured the implanted locations of the recording chambers, enabling the co-registration of .5mm isotropic T1 weighted (MPRAGE) magnetic resonance images and anatomical information with the electrode array. A SC96 semi-chronic microdrive array (Gray Matter Research, USA) was inserted into the recording chambers and sealed for the duration of the recordings. Before recordings were made during the task behavior, about 10-20 electrodes per day were lowered out of the drive to a position past the dura mater where neural activity could initially be observed. Experimental recordings started from these initial positions and on any recording day up to 20 electrodes were moved in small (15-60μm) increments in a random and distributed pattern to allow for continuous sampling of spiking activity during the recordings. Electrode movement was stopped when spiking activity was encountered and spiking activity was not screened for particular functional profiles before the start of the recordings on each day, leading to unbiased sampling of spiking activity across the array.

### Experimental hardware and data acquisition

Eye positions were continuously monitored with an infrared eye tracking system at 120 Hz (ISCAN, USA) and reach touches were registered using an acoustic touch screen (ELO Touch Systems, USA). Both signals were digitized at 1 kHz. Visual stimuli were presented on a LCD monitor (Dell) placed at about 40cm from the subjects’ eyes immediately behind the touch screen. The experimental environment was controlled with custom written LabView (National Instruments) software on a real-time embedded system (NI PXI-9194, National Instruments). Behavioral events were synchronized with neural recordings using photodiode signals on the bottom corner of the LCD monitor. Fluid rewards were delivered with an electronically controlled solenoid.

The microdrive arrays were equipped with glass coated tungsten electrodes (Alpha-Omega, .7-1.2 MΩ impedance at 1 kHz) of variable length conforming to the individual customization of the microdrives and allowing for an overall travel distance (throw) of 2 cm per electrode. The drives contained 96 electrodes spaced at 1.5mm and allowed for individual, bidirectional manual control of electrode movement along one cardinal axis with an approximate maximal resolution of 15μm. During the neural recordings signals were amplified, low-pass filtered at 6 kHz and digitized at 30 kHz using 16 bit resolution with the lowest significant bit equal to 0.1μV (NSpike NDAQ System, Harvard Instrumentation Lab, USA; x10 gain headstage, Multichannel Systems, Germany) and streamed to disc (custom C and MATLAB code). Recordings were referenced to separately implanted ground screws at centimeter distances from the drives that were in contact with the dura mater.

From the raw signals we obtained multi-unit activity offline by high-pass filtering the data at 300 Hz and thresholding with a median-based robust threshold estimate of 3.5 standard deviations below signal mean. Single unit activity was obtained by performing principal component analysis on the multi-unit spike waveforms, following by over-clustering of the first three dimensions by k-means and cluster merging through visual inspection using custom code written in MATLAB (The Mathworks, USA). Unit clusters were tracked in successive 100s data windows to account for non-stationarity in the recordings. All multi and single unit activity was visually inspected based on activity profiles during the tasks and only entered into the data base for the analyses presented here when neural responses and stability of the recordings was observed.

### Behavioral tasks

Animals were trained to perform movement tasks involving reaching while maintaining fixation as well as making saccades while maintaining central touch on a touch screen. On every recording day, animals first performed variable amounts of simple delayed center-out movements to single targets arranged on a circle at a radial distance of 10 degree visual angle. This data is not further discussed here. After the center-out movement tasks the animals performed the dynamic foraging task as detailed in the main text (**Fig 1a**). The choice targets were located at an eccentricity of 10 degree visual angle and were always located on a horizontal axis. Animals had to maintain touch and/or fixation within 2 degree visual angle of all targets for correct task performance. Upon arriving at the target of choice, animals had to hold their gaze or touch at the location for at least 300 ms before fluid rewards were delivered. Successive trials were interrupted by a randomized inter trial interval of 800 – 1300 ms. Three isoluminant colors were used to instruct the animals about movements. The central target was yellow signaling the animals to acquire touch as well as fixation. The eccentric movement targets were either green or red signaling the animals that either reach and fixate or saccade and touch movements were required respectively. As a go cue the central target dimmed into a darker gray that for clear visibility was less luminant than the colors used for instructing the movements. The reach and saccade conditions alternated in longer blocks of random length on every day (Monkey J: Reaches, mean 102 trials, SD 86 trials; Saccades, mean 127 trials, SD 78 trials; Monkey H: Reaches, mean 164 trials, SD 96 trials; Saccades, mean 186 trials, SD 73 trials). Block transitions were unsignaled (apart from the change in target color) and the starting condition was chosen randomly on every day. Both animals performed at least one block of reaches and one block of saccades per recording session. Overall monkey J performed the task on 62 neural recording sessions for a total of 157 reach and 148 saccade blocks and a total of 34994 trials. Monkey H performed the task on 45 neural recording sessions for a total of 109 reach and 102 saccade blocks and a total of 36949 trials. Monkey J reached with the left arm and neural recordings were made in the right hemisphere. Monkey H reached with the right arm and neural recordings were made in the left hemisphere.

### Reward Drifts and objectively optimal values

During the dynamic choice task, the reward magnitudes at the two movement target locations drifted in an uncorrelated way according to gaussian random walks. The design of the reward drifts and the Kalman filter predictions were based on work on human reinforcement learning systems^15^. Briefly, we measured the opening times of the solenoid valve for which we could reliably and linearly deliver fluid volumes. Based on these measurements on trial *t* the reward at target *m* was specified as an opening time of the valve between 220 ms and 400 ms and was drawn from a Gaussian distribution around a mean μ_t,m_ with standard deviation σ_0_ 15.5 ms. On every trial the mean drifted with:

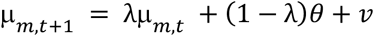

where the decay parameter *λ* was 0.9836, the decay center θ was 310 ms and the zero mean Gaussian diffusion noise *v* had a standard deviation of σ_n_ = 11.16 ms.

We generated 24 pairs of reward drifts for the choice task. In order to coarsely balance the statistics of the drifts across reaches and saccades as well as the two target locations, we replayed the drifts after several days. We either replayed the same drifts for days on which the animals started out with saccades and had sampled the drift starting with reaches already (or vice versa), or we switched the association of spatial movement target and drift.

To objectively predict value states of the movement targets we used a Kalman filter to track the reward drifts. If movement target *m* was chosen the predicted mean of the drift was updated according to:

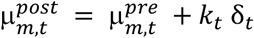

Where *δ*_t_ is the prediction error given as the difference between received reward r_t_ and its prediction 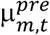 and k_t_ is the learning rate:

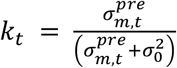

Together with the mean the predicted variance of the chosen option was updated according to:

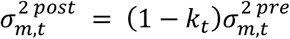

The mean and variance of the unchosen movement target were unaffected and predictions for the next trial for all *m* were obtained as:

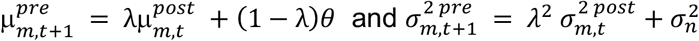

We used the predicted means from this tracking process as the objectively optimal value state predictions on any given trial.

### Behavioral modeling

We used nested model testing based on logistic regressions to investigate the choice behavior of the animals in detail. Specifically, we hierarchically fit all parameter of the model:

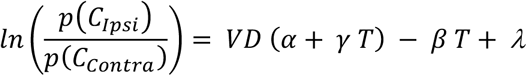

Where C represents the choices, VD the difference in objective values at the two targets and T the behavioral timing on any given trial. We compared the models using the Akaike information criterion (AIC) and interpreted AIC differences of 10 as strong evidence for a better fit to the data^29^. We chose the best models for behavior (**see Fig S1, S2**) based on the AIC as well as the observation that all regressors of the models were a significant (p < 0.05, **see Table S1**). We combined trials across recordings sessions to obtain the model fits.

### Neural data analyses

We used Linear Discriminant Analysis to classify trials according to movement conditions based on the spiking activity of simultaneously recorded units in frontal cortex^30^. We made binary classifications of the trials either based on the direction of the movements that animals made (ipsilateral vs. contralateral) or the temporal condition of the movements (immediate vs. delayed). We refer to these classification performances as choice and enaction information, respectively. Choice information was computed separately for immediate and delayed trials and all trials were pooled to compute the enaction information. The classifiers were trained based on the single trial activity of simultaneously recorded units (range 1 - 40) within sliding 50 ms windows stepping through time in 10 ms steps. In all multivariate analyses the identical units and ensembles contributed to the classification for reaches and saccades. For all multivariate and univariate analyses within the anatomical subdivisions we included all units recorded in the respective subdivision (**Fig 3, S3, S4, S5, S6, S7**). For all multivariate analyses pooling units into array-wide ensembles (**Fig 4, 6, S8, S9**) we only included ensembles with at least 10 units from which at least 1 unit was present in each anatomical subdivision (**see Fig S5 a, b**). We excluded units from the array wide ensembles if their inclusion would have reduced the number of trials for that ensemble by 100 e.g. due to loss of unit isolation. Classification was only performed if at least 5 trials of each movement condition were available and each trial was held out for training when classified, training only on all other trials (leave-one-out cross validation). For the classification results presented in main text figure 3 we only included data of each trial up to 50 ms before its measured reaction time (end of central touch or fixation). We assessed classification performance as the difference in true positive and false positive rates of the classifiers to account for eventual differences in trial numbers between the conditions. To test for significant classification performance, we performed random permutation tests. We obtained resample distributions of classification performances under the null hypothesis by randomly flipping the sign of the classification performances of individual ensembles 10,000 times. We then used these resample distributions to non-parametrically compute p-values. Where appropriate we corrected for multiple comparisons by controlling the false discovery rate^31^.

We estimated the latencies of the enaction information as the timepoint at which the information reached half maximum within 200 – 400 ms after target onset. We only estimated latencies for an anatomical subdivision if significant classification performance was present. We obtained confidence estimates of the latencies by bootstrapping.

To regress single trial value differences against the choice classification performance we obtained single trial classification probabilities (posterior class probabilities) according to the procedures described above for average spiking activity within 500 ms windows centered at 4 non-overlapping time points of interest during delayed choice trials. We defined the baseline time-point as 250 ms before the target onset, the commit time-point as the local peak in classification performance within the first 500 ms during delayed choices (i.e. animal and movement specific), the pre-go time point as 250 ms before the go cue on delayed choices and the reaction time time-point as the median reaction time on delayed choices (i.e. animal and movement specific). We regressed the value differences between the two targets and the classification probabilities separately for ipsilateral and contralateral movements using Pearson correlations. To assess the significance of the regressions we performed random permutation tests by shuffling the trial association of value differences and classifier probabilities 10,000 times. We then squared the regression coefficients as well as the resample distribution and non-parametrically converted the data to the corresponding z-scores and averaged the z-scores for each ensemble across ipsilateral and contralateral movements. We then tested whether the average z-scores across ensembles were significantly positive using t-tests.

For the cross-classification analyses the classifiers were trained separately on either all immediate choice or all delayed choice trials to predict either delayed or immediate choices, respectively. We obtained corrected cross-classification performance by subtracting the product of the classification performance on delayed and immediate choices for each specific time-point combination of every classification. We assessed significance of the cross-classification performances with permutation tests as described for the within condition classifications above. To assess the dynamics of how action plans are represented across the frontal cortical ensembles we split all neurons into two pools encoding choices towards the ipsilateral or contralateral side, respectively. For this analysis we selected all units that exhibited a significant firing rate difference between ipsilateral and contralateral choices within a 500ms time window before the reaction time. After that we split neurons into pools encoding for the ipsilateral and contralateral movement direction according to the area under the curve of a receiver operating characteristic (AUROC), computed for every unit with ipsilateral movements as the consistent positive class. Units with values greater than 0.5 were treated as the ipsilateral encoding pool and units with values less than 0.5 were treated as the contralateral encoding pool. Before averaging the activity of the units during ipsilateral and contralateral choices we normalized the firing rate of each unit by dividing through the maximum firing rate across all conditions (ipsi- and contralateral movements as well as immediate and delayed trials). Error bars throughout all analyses represent 95% confidence intervals obtained by bootstrapping. Analyses were performed using MATLAB (The Mathworks, USA).

## Supplementary results

### Behavioral modelling of context dependent decision making

To assess how choices were revised in real-time, we modeled the temporal evolution of choice computations from target presentation to the movement choice (**Fig S1, S2**). The best fitting IR model suffered a significant spatial bias (p < 0.05, **Table S1, Fig. S1 a, b, bottom row**). In addition, the way that the animals allocated their IR choices also depended on behavioral timing (p < 0.05, **Table S1, Fig. S1 a,b**) As reaction times got longer the animals allocated their IR choices more evenly across the two movement targets. In other words, fast reaches exhibited the strongest ipsilateral bias while slower reaches sampled the targets more evenly.

The DR exhibited a pattern in which the animals allocated their choices in a more balanced way. As a result, the animals recovered hits on the contralateral side bringing them up to comparable levels with the ipsilateral side (**Fig 2 a, Fig. S1 c, d**). Strikingly, the timing parameters (reaction time, elapsed time) influenced the allocation of choices in an opposite pattern between IR and DR. That is, the fastest DR reaction times were associated with a contralateral bias that then disappeared for longer reaction times. This effect manifested in opposite signs of the model weights for the reaction time regressors (**Table S1**).

Overall, animals made qualitatively different reach decisions depending on the temporal context. While IR were suboptimal and biased, choices became gradually more evenly distributed across the two movement options, achieving an objectively more optimal income for DR. Strikingly, the transition from early suboptimal, to later more optimal choices was associated with an inversion of a bias that depended on behavioral timing. Thus, the animals dynamically changed their preferences towards a greater impact of objectively optimal values during DR and this change was linked to dynamics in sensorimotor circuits.

The animals engaged in the identical task also with saccades. Saccades are fast and energetically less costly. We predicted that the different dynamics of the oculomotor system may lead to a different pattern of decision making under otherwise identical conditions. Specifically, for saccades the decisions should be less influenced by particular actions themselves and reflect a selection process more closely tied to the objective values of the targets. In line with this prediction, saccade choices of the animals were more stable and optimal than the reach choices (**Fig. S2**).

For monkey J, IS and DS had a comparable pattern (**Fig. 3a**) of hits with a symmetrical appearance of hits for both spatial locations (**Fig. S2c**). The IS choices had a small reaction time dependent bias similar to that of reaches (p < 0.05.) with a bias towards the ipsilateral side for very fast reaction times that disappeared for longer reaction times (**Fig. S2a**). The DS of monkey J were then strongly driven by the difference in objective values and independent of reaction times (**see Table S1, Fig. S2a**).

Monkey H exhibited a comparable foraging pattern with symmetrical hits for IS and DS (**Fig. 3b**). However, for monkey H both IS and DS were best modelled with reaction time dependent biases (**see Table S1, Fig. S2b**). Interestingly, for monkey H the saccade choices exhibited an inversion of a reaction time dependent bias, similar to what we observed for reaches. A contralateral bias increased with reaction for IS while for DS the contralateral bias decreased with time. The increasing bias for IS choices led to a reaction time dependent decrease in hits on the ipsilateral side (**Fig. S2d**). That is, with increasing reaction time monkey H’s choices were increasingly biased towards the contralateral side. Both monkeys’ saccade choices had a slight overall contralateral bias leading to an asymmetry in the hits for the ipsi- and contralateral sides for small value differences (**Fig. S2c, d**).

Overall the choices of the animals were strongly influenced by the temporal and action contexts of the decisions. Most strikingly, reach choices were associated with a dynamic change in choice preferences of the animals, with qualitatively different decisions made immediately or after a brief delay. These behavioral observations are consistent with the idea that the context dependent choices may have resulted from an ongoing deliberation that was dynamically read out by the process of enaction to support current preferences.

### Decision information in array wide neuronal ensembles

We pooled simultaneously recorded units into array wide ensembles to capture the distributed decision information across all recorded areas (**Fig S5**). Pooling alters the multivariate composition of the ensembles and may lead to different dynamics of information resulting from our classification approach. To reveal the patterns of array wide information, we decoded choice and enaction information (**Fig S5c, d**). Pooled ensembles captured highly similar dynamical features a compared to the ensembles within individual anatomical subdivisions (**Fig 3, S3, S4**). Choice information was significantly present during the baseline interval (p<0.05, FDR corrected, **Fig S5c, d**). Choice information exhibited a brisk increase shortly after the onset of the choice targets and then further increased for immediate choices or slightly dipped and then plateaued for delayed choices. The enaction information also exhibited comparable dynamics and increased sharply at about the time when choice information started to diverge between immediate and delayed choices.

## Supplementary Table S1 : Nested model testing of choice behavior

**Abbreviations**
IR: immediate reach

DR: delayed reach

RT: reaction time

DL: delay

IS: immediate saccade

DS: delayed saccade

AIC: Akaike information criterion

The full model containing all parameters was fit as:

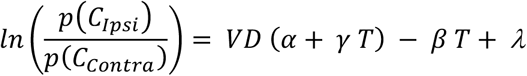

Where C represents choices, VD the difference in objective values at the two options and T the behavioral timing on any given trial. RT is the reaction time, while ET is the elapsed time since target presentation obtained as the delay plus the reaction time.

| Monkey J | IR, Timing = RT |  | DR, Timing = ET |  |
| --- | --- | --- | --- | --- |
|  | AIC | Param. | AIC | Param. |
| constant ($\lambda$) | 6,680 | 0.5 | 12,947 | -0.1 |
| $\alpha$ | 6,498 | 1.71 | 11,486 | 2.23 |
| $\beta$ | 6,843 | 0.68 | 12,967 | 0.09 |
| constant, $\alpha$ | 6,145 | 0.57, 1.85 | 11,453 | -0.13, 2.25 |
| constant, $\beta$ | 6,609 | 1.03, 1.10 | 12,934 | -2.4, -0.3 |
| constant, $\alpha$, $\beta$ | 6,084 | 1.08, 1.85, 1.08 | **11,441** | -0.28, 2.25, -0.31 |
| constant, $\alpha$, $\beta$, $\gamma$ | **6,069** | 1.14, 2.60, 1.19, -1.58 | 11,443 | -0.28, 2.23, -0.31, 0.03 |
| **Chosen Model**, Parameter $p < 0.05$, Parameter $p > 0.05$ | | | | |

| Monkey H | IR, Timing = RT |  | DR, Timing = ET |  |
| --- | --- | --- | --- | --- |
|  | AIC | Param. | AIC | Param. |
| constant ($\lambda$) | 5,805 | 0.97 | 15,311 | 0.26 |
| $\alpha$ | 5,862 | 2.53 | 12,133 | 3.35 |
| $\beta$ | 6,376 | 1.64 | 14,631 | -1.06 |
| constant, $\alpha$ | 4,519 | 1.33, 3.43 | 11,739 | 0.45, 3.55 |
| constant, $\beta$ | 5,691 | 1.52, 1.53 | 13,735 | -1.38, -3.47 |
| constant, $\alpha$, $\beta$ | **4,453** | 1.82, 3.40, 1.36 | **10,571** | -1.22, 3.56, -3.51 |
| constant, $\alpha$, $\beta$, $\gamma$ | 4,455 | 1.83, 3.48, 1.40, -0.21 | 10,573 | -1.21, 3.63, -3.5, -0.15 |
| **Chosen Model**, Parameter $p < 0.05$, Parameter $p > 0.05$ | | | | |

| Monkey J | IS, Timing = RT |  | DS, Timing = ET |  |
| --- | --- | --- | --- | --- |
|  | AIC | Param. | AIC | Param. |
| constant ($\lambda$) | 8,353 | -0.006 | 15,450 | 0.02 |
| $\alpha$ | 4,941 | 5.50 | 8,461 | 6.41 |
| $\beta$ | 8,272 | 0.46 | 15,488 | -0.06 |
| constant, $\alpha$ | 4,941 | 0.05, 5.50 | **8,454** | 0.82, 6.41 |
| constant, $\beta$ | 7,976 | 0.96, 2.16 | 15,450 | -0.02, -0.09 |
| constant, $\alpha$, $\beta$ | **4,790** | 0.89, 5.43, 1.87 | 8,455 | 0.03, 6.41, -0.09 |
| constant, $\alpha$, $\beta$, $\gamma$ | 4,786 | 0.90, 6.08, 1.88, -1.44 | 8,454 | 0.03, 6.06, -0.09, 0.73 |
| **Chosen Model**, Parameter $p < 0.05$, Parameter $p > 0.05$ | | | | |

| Monkey H | IS, Timing = RT |  | DS, Timing = ET |  |
| --- | --- | --- | --- | --- |
|  | AIC | Param. | AIC | Param. |
| constant ($\lambda$) | 8,457 | -0.4 | 14,898 | -0.23 |
| $\alpha$ | 5,987 | 4.45 | 10,505 | 4.34 |
| $\beta$ | 8,396 | 1.43 | 15,026 | 0.17 |
| constant, $\alpha$ | 5,471 | -0.76, 4.86 | 10,118 | -0.49, 4.58 |
| constant, $\beta$ | 8,390 | -0.13, 1.10 | 14,759 | -0.63, -0.9 |
| constant, $\alpha$, $\beta$ | 5,390 | -0.30, 4.91, 1.54 | 10,088 | -0.73, 4.55, -0.55 |
| constant, $\alpha$, $\beta$, $\gamma$ | **5,321** | -0.49, 6.47, 0.85, -4.93 | **10,078** | -0.77, 5.09, -0.64, -1.18 |
| **Chosen Model**, Parameter $p < 0.05$, Parameter $p > 0.05$ | | | | |

## Supplementary Figure Legends

**Figure S1:**
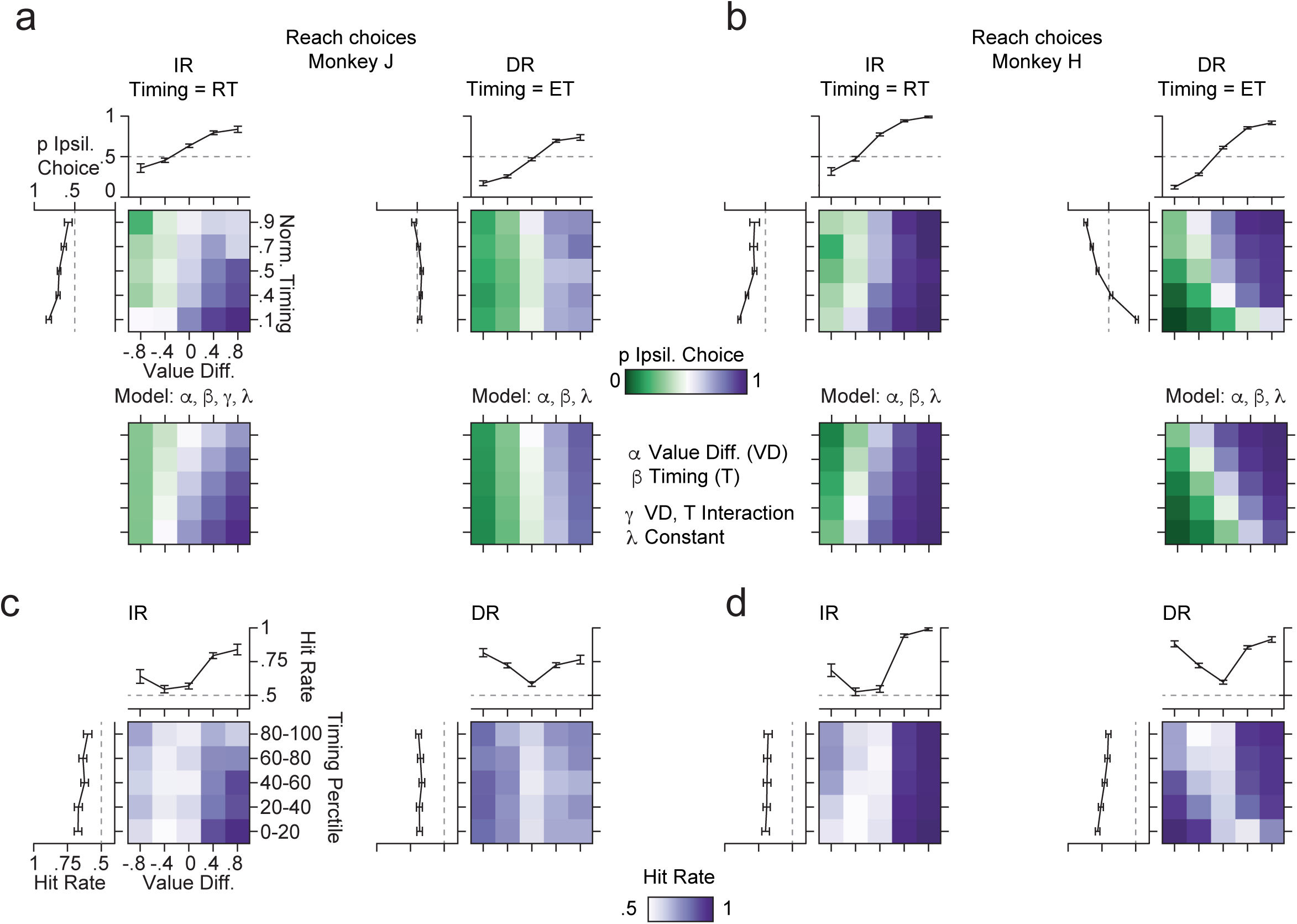
Choice allocation, behavioral modelling and foraging success for reach choices. a) Empirical choices and best model predictions for reach decisions of monkey J. Immediate reach (IR) and delayed reach (DR) choices are shown as a function of behavioral timing and the differences of the objectively optimal values at the two target locations. Positive value differences indicate a higher value at the ipsilateral target, negative value differences indicate a higher value at the contralateral target. IR are shown with reaction time (RT) as timing parameter. DR are shown with elapsed time (ET) since target onset as timing parameter, representing the sum of the delay and RT. The line plots show the choice probabilities based on the two factors separately. All error bars depict 95% confidence intervals. The best models were chosen according to a nested model selection (**see Table S1**) and the contributing parameters are shown above the model prediction plots. α, value difference; β, timing; *γ*, multiplicative interaction of the value difference and the timing; *λ*, constant term. b) Same as a but for monkey H. c) Foraging success of monkey J during reach choices. The hit rate is the probability of the animal choosing the objectively higher valued target. Hit rate is shown for the value and timing factors as well as other conventions described above. d) same as c for monkey H.

**Figure S2:**
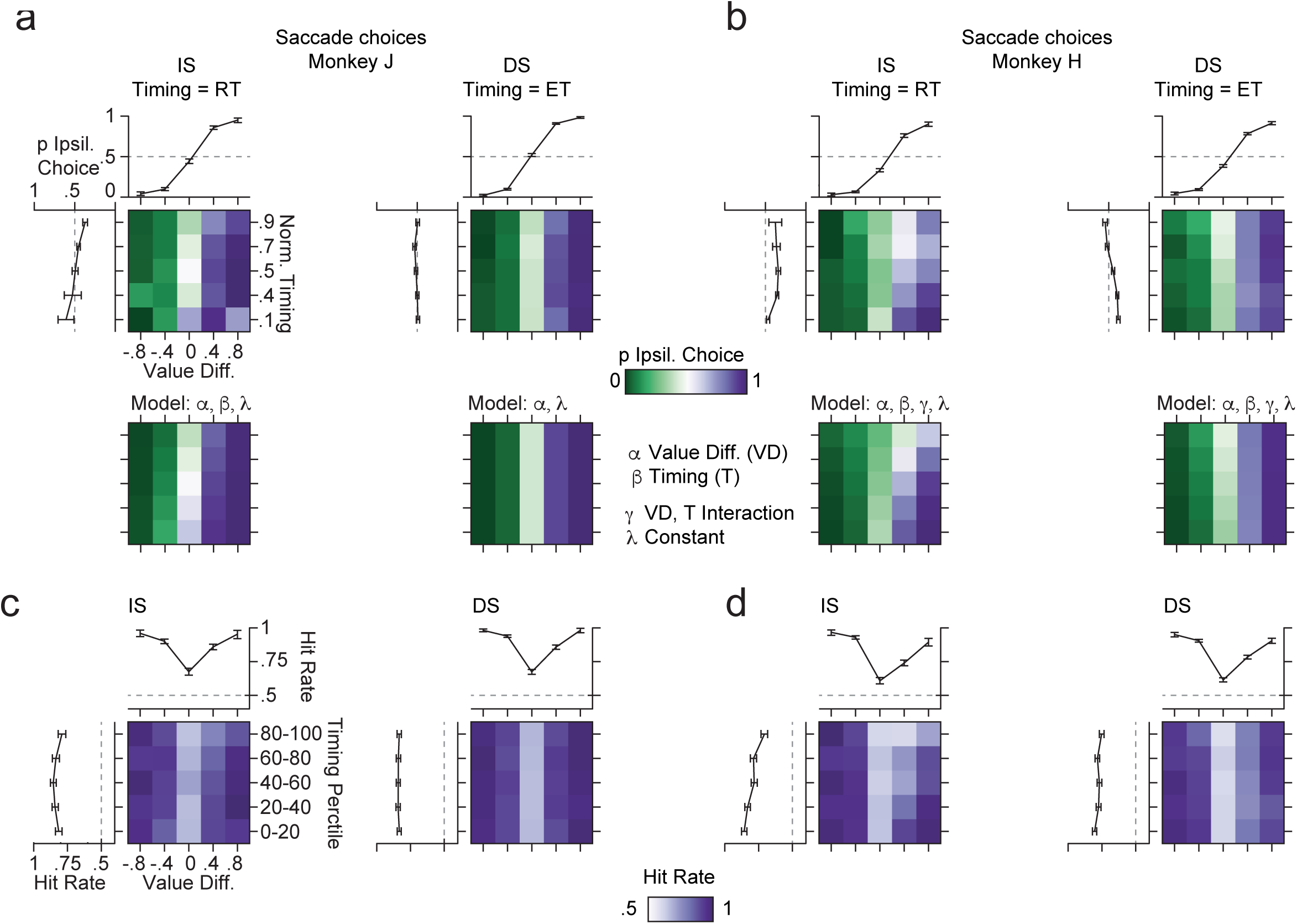
Choice allocation, behavioral modelling and foraging success for saccade choices. Same as Fig S1 for saccade choices.

**Figure S3:**
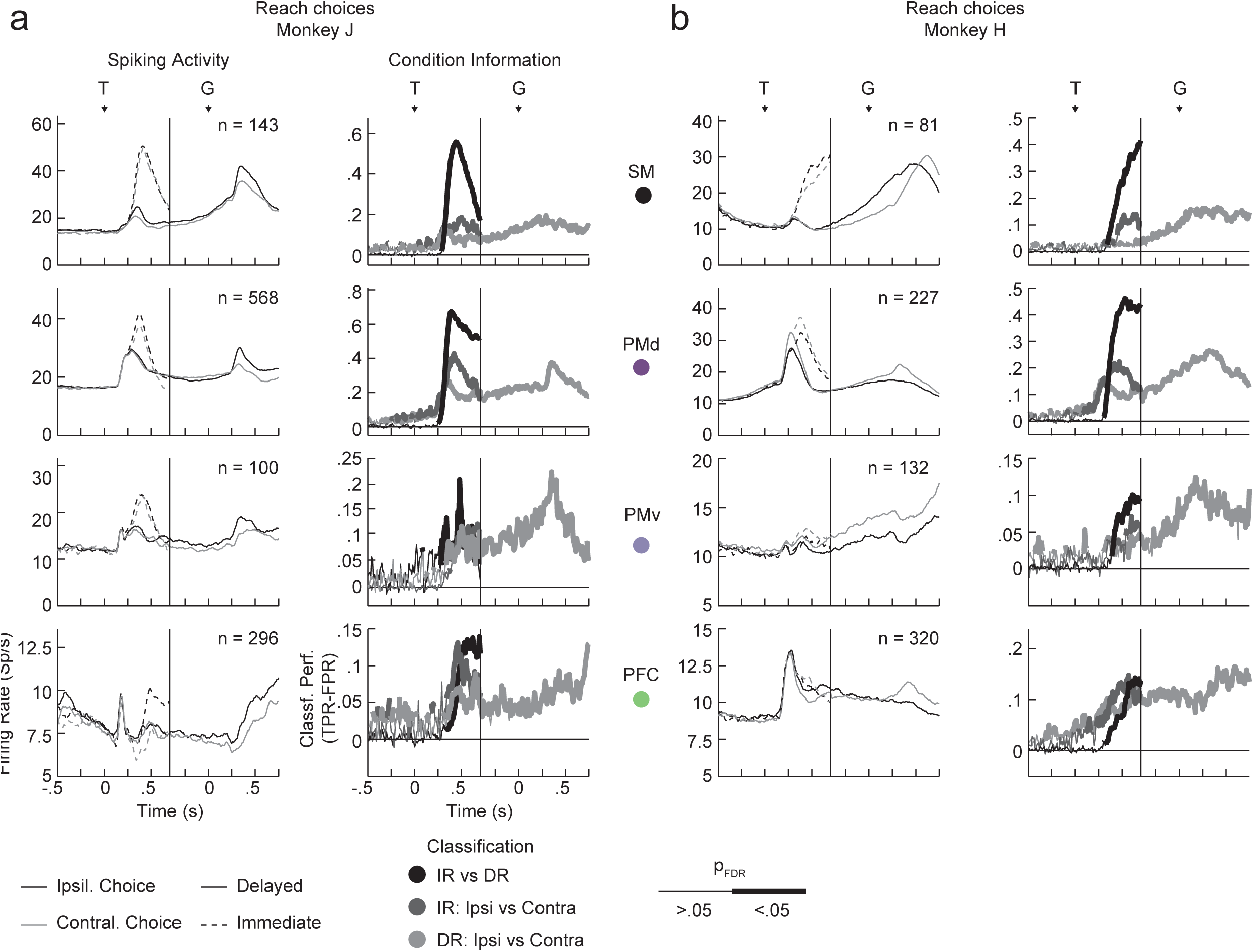
Spiking activity and classification performance within anatomical subdivisions during reach and fixate decisions. a) The left column shows the average spiking activity during the four principal conditions (ipsilateral and contralateral choices as well as immediate and delayed choices) of units within 4 anatomical subdivisions covered by the electrode arrays. The number of contributing units is shown within each panel. T, target onset; G, go cue. The right column shows the classification performance of three classifiers based on the spiking activity. We revealed enaction information as the classification of immediate reaches (IR) vs delayed reaches (DR) and choice information for the IR and DR separately by classifying trials into ipsi- and contralateral movement choices. The classification performances are plotted as thick lines where significant classification performance was detected (p< 0.05, FDR corrected, permutation tests). SM, somato-motor; PMd, dorsal premotor cortex; PMv, ventral premotor cortex; PFC, prefrontal cortex. b) Same as A for monkey H.

**Figure S4:**
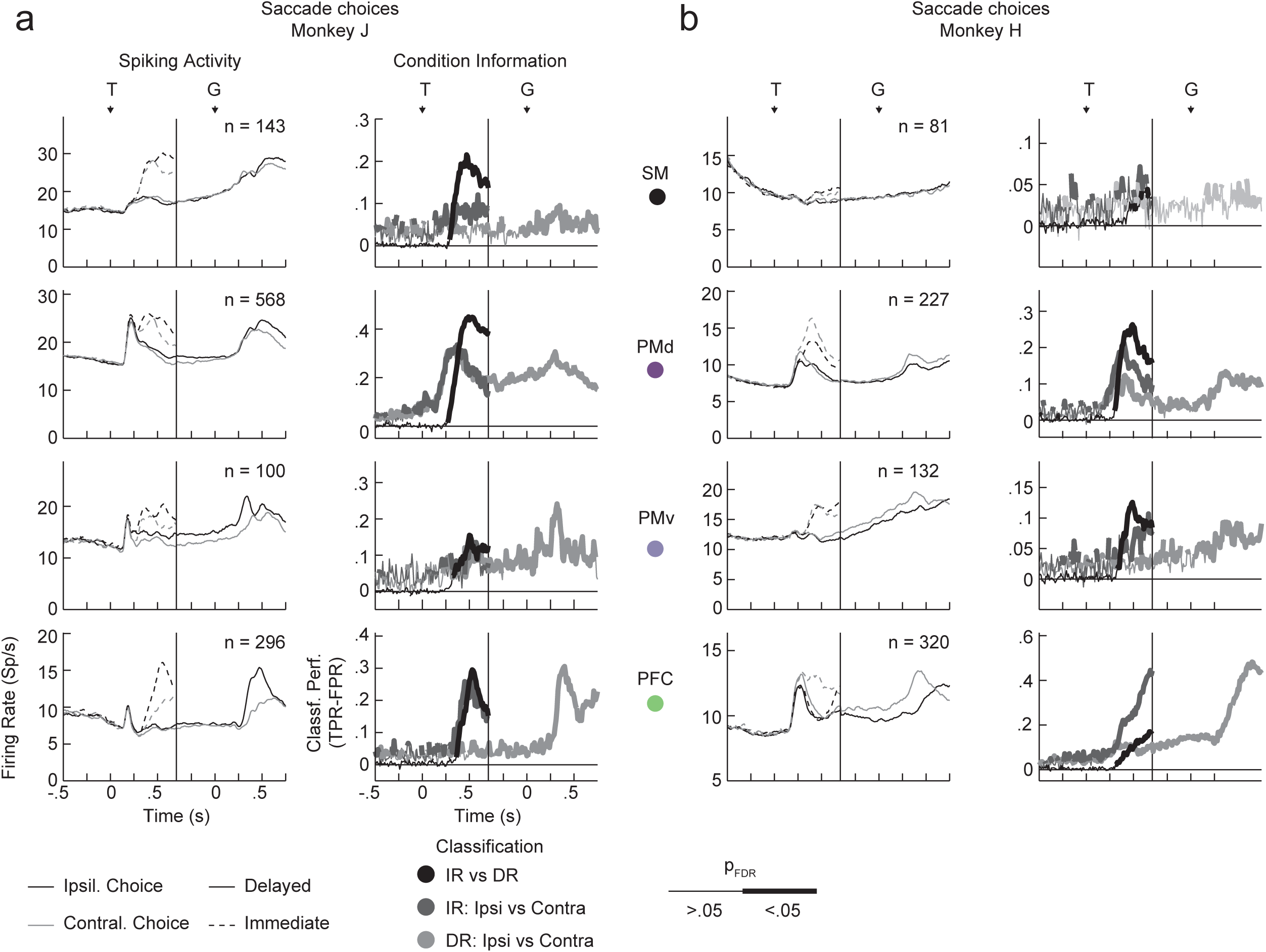
Spiking activity and classification performance within anatomical subdivisions during saccade and touch decisions. Same as **Fig S3** but for saccade and touch choices.

**Figure S5:**
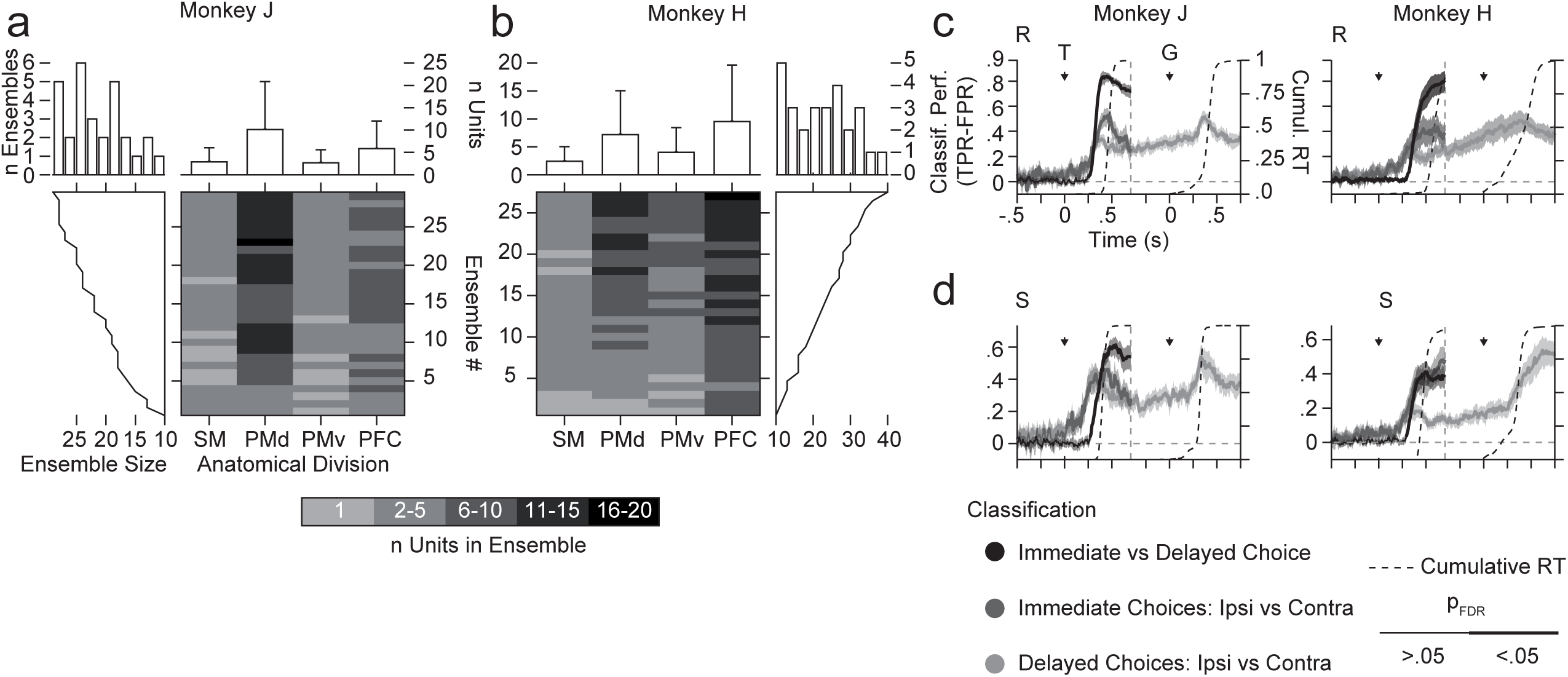
Array wide ensemble classification. a) Array wide ensemble composition for monkey J. The image shows the number of units from each anatomical subdivision. Ensembles are sorted by their size as seen in the marginal plot at the side of the image. The average number of units per ensemble is shown on top of the image plot and a histogram of ensemble sizes is given at the top left. b) same as a for monkey H. c) Classification performances for reach decisions for monkey J and monkey H. We computed enaction information as the classification performance between immediate and delayed choices and choice information separately for immediate and delayed choices as the classification performance between ipsi- and contralateral movements. Significant classification performance is shown as thick lines (p < 0.05, FDR corrected, permutation tests). Shadings depict 95% confidence intervals. TPR, true positive rate; FPR, false positive rate; T, target onset; G, go cue. The cumulative distribution of reaction times (RT) for immediate and delayed choices is shown as dotted lines. d) Same as c but for saccade and touch data. Note that the same ensembles contributed to the classification of reach and saccade data.

**Figure S6:**
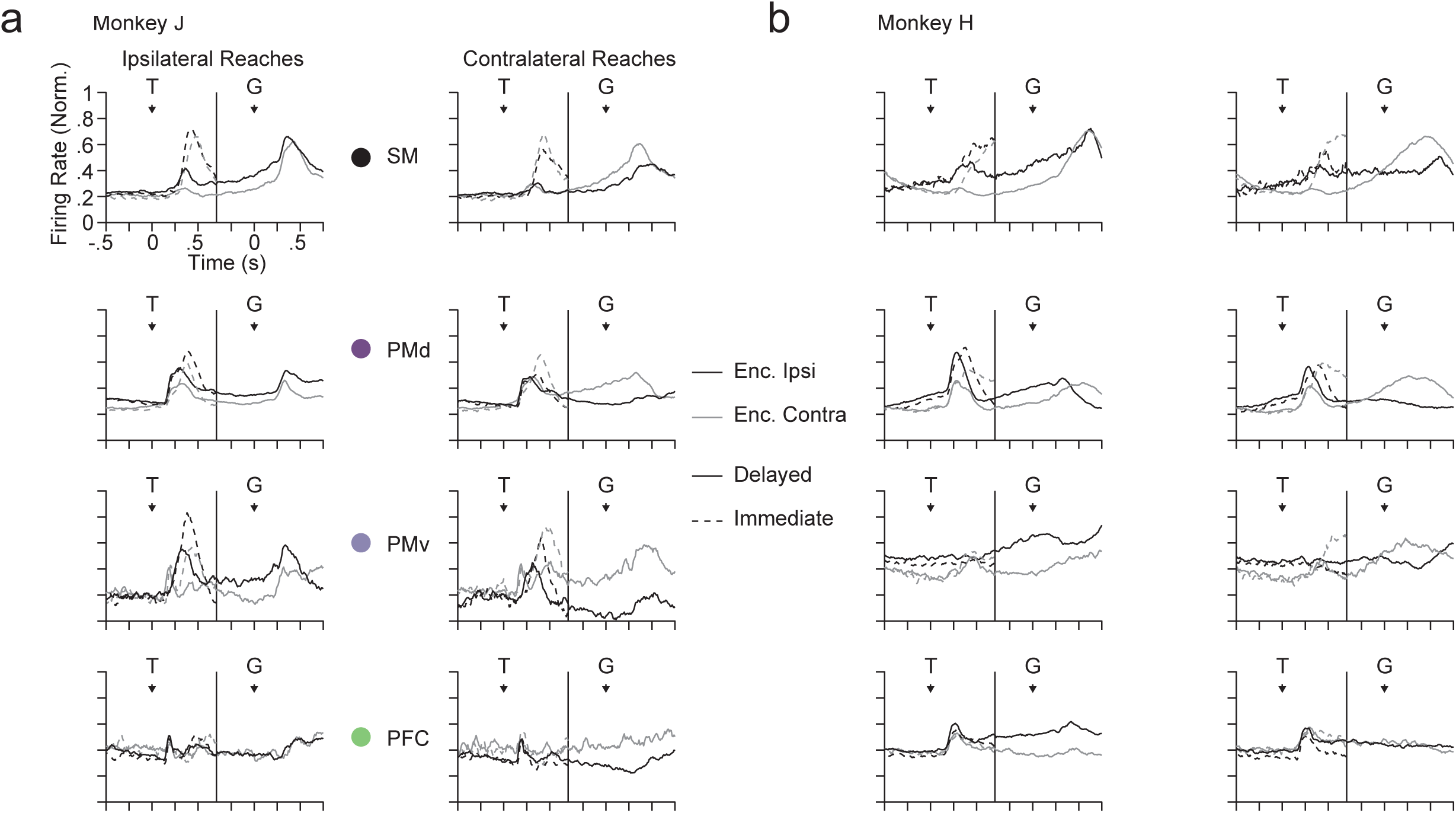
Sensorimotor circuits track action preferences. Same conventions as **Fig 5** but for individual anatomical subdivisions during reach and fixate choices. a) Activity profiles for ipsilateral (left column) and contralateral (right column) reaches for pools of neurons encoding the ipsilateral (black) and contralateral (gray) during delayed (solid lines) and immediate (dotted lines) choices. Each row depicts activity in one anatomical subdivision. See **Fig 5** for the number of neurons found to encode choices in each anatomical subdivision. SM, somato-motor; PMd, dorsal premotor cortex; PMv, ventral premotor cortex; PFC, prefrontal cortex. T, target onset; G, go cue; b) Same as A for monkey H.

**Figure S7:**
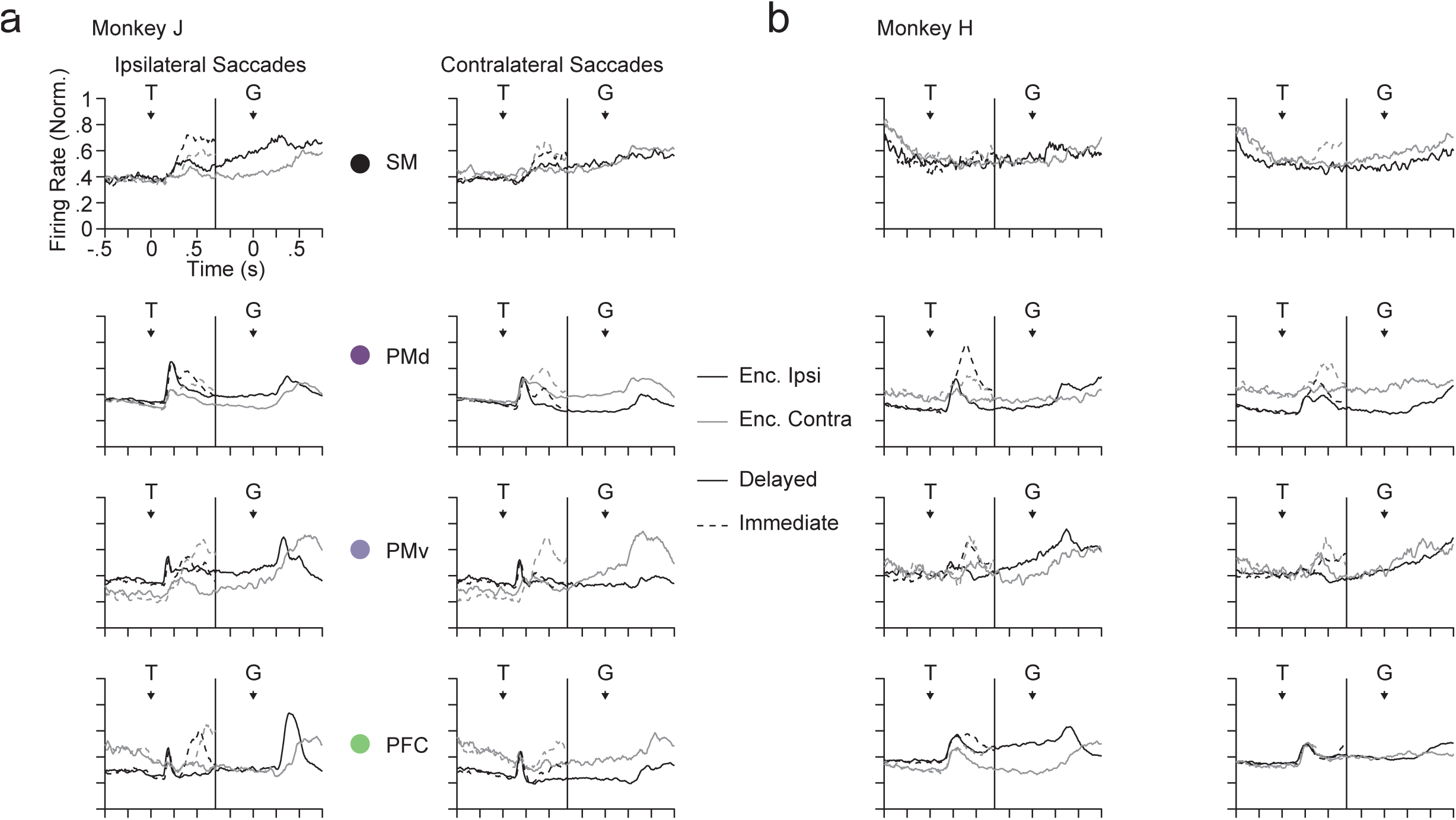
Same as Fig S6 for saccade and touch data.

**Figure S8:**
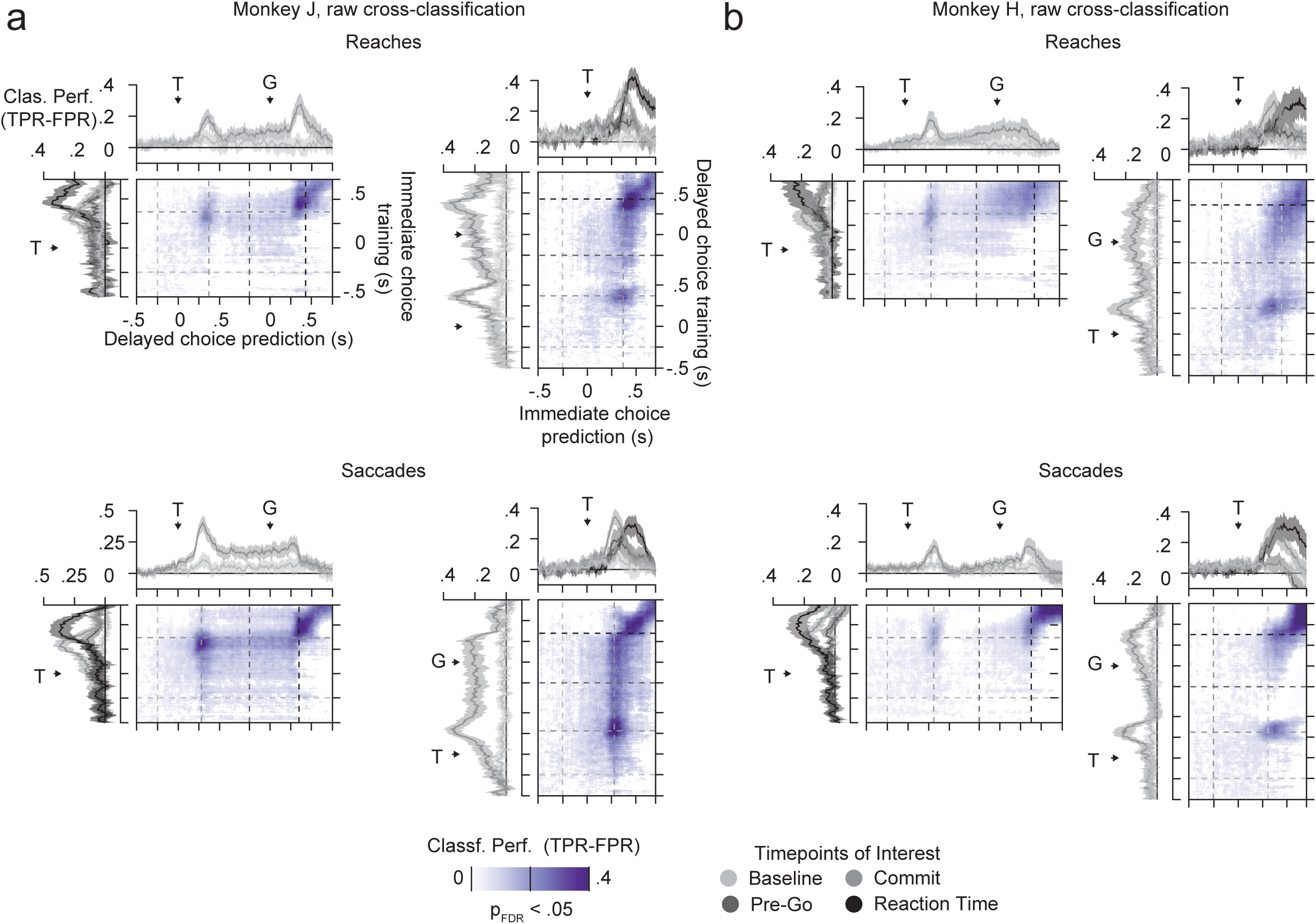
Raw cross-classification a) Classification performance of training classifiers on different time points of immediate choices and predicting delayed choices (left) or vice versa (right). The image plots show significant cross-classification as purple (p<0.05, FDR corrected). The line plots on the sides show slices of the image plot along with 95% confidence intervals (shadings) along the four time-points of interest during the trials as used previoulsy: baseline, commit, pre-go and reaction time. The upper row depicts cross-classification of reaching, the bottom row shows cross-classification of saccades. b) Same as a for monkey H.

**Figure S9:**
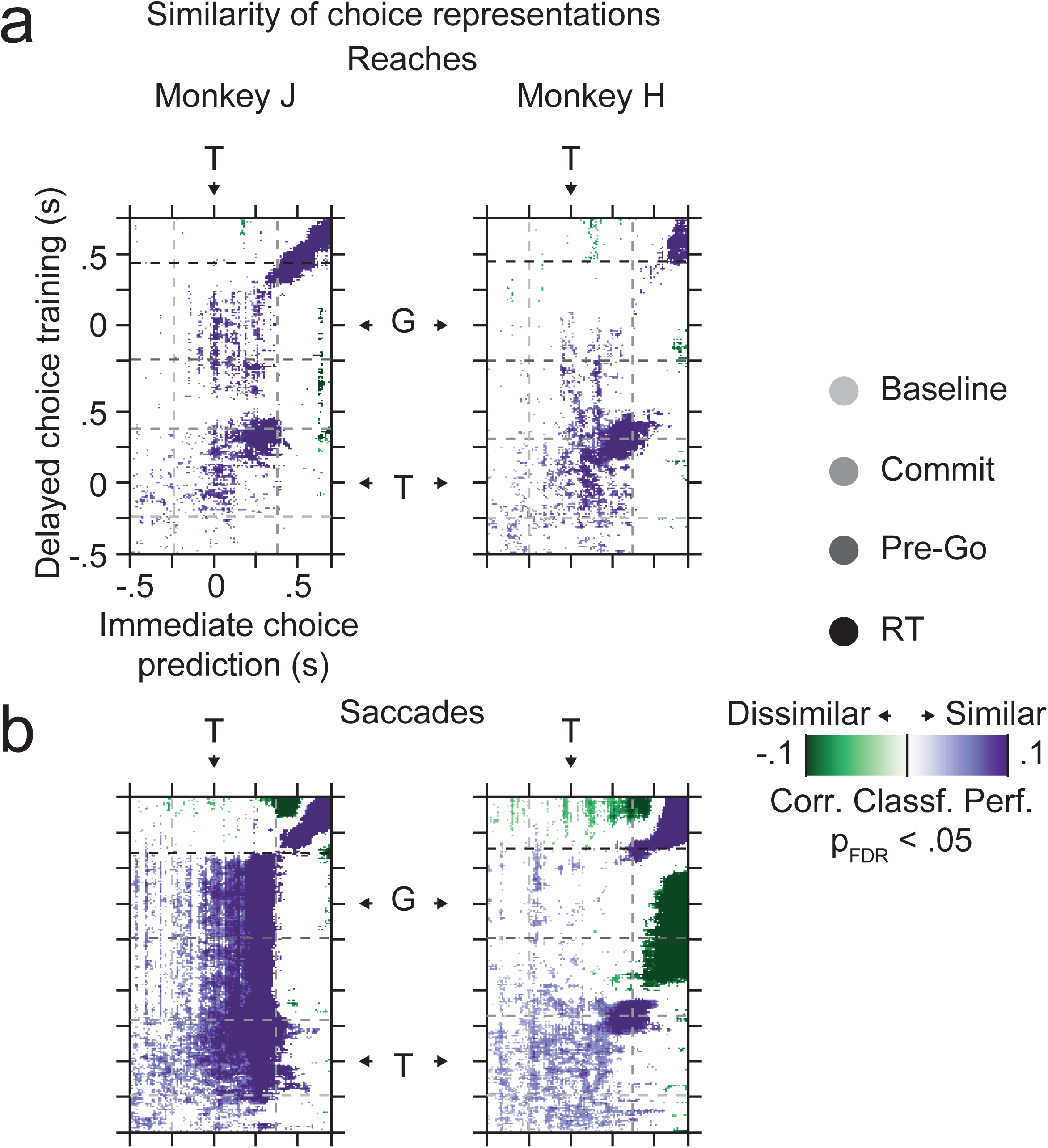
Similarity of the choice representations obtained through corrected cross-classification for training on delayed choices and predicting immediate choices. Same conventions as **Fig 6**.

## References

1. Padoa-Schioppa, C. & Conen, K. E. Orbitofrontal Cortex: A Neural Circuit for Economic Decisions. Neuron 96, 736–754 (2017).

2. Cisek, P. Making decisions through a distributed consensus. Curr. Opin. Neurobiol. 22, 927–936 (2012).

3. O’Donoghue, T. & Rabin, M. Doing it now or later. Am. Econ. Rev. 89, 103–124 (1999).

4. Thaler, R. H. & Shefrin, H. An economic theory of self-contol. Journal of Political Economy 89, 392–406 (1981).

5. Loewenstein, G. Out of Control: Visceral Influence on Behavior. Organ. Behav. Hum. Decis. Process. 65, 272–292 (1996).

6. Resulaj, A., Kiani, R., Wolpert, D. M. & Shadlen, M. N. Changes of mind in decisionmaking. Nature 461, 263–266 (2009).

7. Kahneman, D. Maps of Bounded Rationality: Psychology for Behavioral Economics. Am. Econ. Rev. 93, 1449–1475 (2003).

8. Cos, I., Belanger, N. & Cisek, P. The influence of predicted arm biomechanics on decision making. J. Neurophysiol. 105, 3022–3033 (2011).

9. Roitman, J. D. & Shadlen, M. N. Response of neurons in the lateral intraparietal area during a combined visual discrimination reaction time task. J. Neurosci. 22, 9475–89 (2002).

10. Platt, M. L. & Glimcher, P. W. Neural correlates of decision variables in parietal cortex. Nature 400, 233–238 (1999).

11. Padoa-Schioppa, C. & Assad, J. A. Neurons in the orbitofrontal cortex encode economic value. Nature 441, 223–226 (2006).

12. Pesaran, B., Nelson, M. J. & Andersen, R. A. Free choice activates a decision circuit between frontal and parietal cortex. Nature 453, 406–409 (2008).

13. Cisek, P. & Kalaska, J. F. Neural mechanisms for interacting with a world full of action choices. Annu. Rev. Neurosci. 33, 269–298 (2010).

14. Pesaran, B. Neural correlations, decisions, and actions. Curr. Opin. Neurobiol. 20, 166–171 (2010).

15. Daw, N. D., O’Doherty, J. P., Dayan, P., Seymour, B. & Dolan, R. J. Cortical substrates for exploratory decisions in humans. Nature 441, 876–9 (2006).

16. Cisek, P. Cortical mechanisms of action selection: the affordance competition hypothesis. Philos. Trans. R. Soc. Lond. B. Biol. Sci. 362, 1585–99 (2007).

17. Daw, N. D., Niv, Y. & Dayan, P. Uncertainty-based competition between prefrontal and dorsolateral striatal systems for behavioral control. Nat. Neurosci. 8, 1704–1711 (2005).

18. Thura, D. & Cisek, P. Deliberation and commitment in the premotor and primary motor cortex during dynamic decision making. Neuron 81, 1401–1416 (2014).

19. Thura, D. & Cisek, P. The Basal Ganglia Do Not Select Reach Targets but Control the Urgency of Commitment. Neuron 95, 1160–1170.e5 (2017).

20. Hanes, D. P. & Schall, J. D. Neural control of voluntary movement initiation. Science 274, 427–30 (1996).

21. Rich, E. L. & Wallis, J. D. Decoding subjective decisions from orbitofrontal cortex. Nat. Neurosci. 19, 973–980 (2016).

22. Wong, Y. T., Fabiszak, M. M., Novikov, Y., Daw, N. D. & Pesaran, B. Coherent neuronal ensembles are rapidly recruited when making a look-reach decision. Nat. Neurosci. 19, 327–334 (2016).

23. Hawellek, D. J., Wong, Y. T. & Pesaran, B. Temporal coding of reward-guided choice in the posterior parietal cortex. Proc. Natl. Acad. Sci. U. S. A. 113, 13492–13497 (2016).

24. Siegel, M., Buschman, T. J. & Miller, E. K. Cortical information flow during flexible sensorimotor decisions. Science 348, 1352–1355 (2015).

25. Kable, J. W. & Glimcher, P. W. The neurobiology of decision: consensus and controversy. Neuron 63, 733–45 (2009).

26. Rangel, A., Camerer, C. & Montague, P. R. A framework for studying the neurobiology of value-based decision making. Nat. Rev. Neurosci. 9, 545–556 (2008).

27. Daw, N. D., Gershman, S. J., Seymour, B., Dayan, P. & Dolan, R. J. Model-based influences on humans’ choices and striatal prediction errors. Neuron 69, 1204–1215 (2011).

28. Walton, M. E., Rudebeck, P. H., Bannerman, D. M. & Rushworth, M. F. S. Calculating the cost of acting in frontal cortex. Ann. N. Y. Acad. Sci. 1104, 340–356 (2007).

29. Burnham, K. P. & Anderson, D. R. Model Selection and Multimodel Inference - A Practical Information-Theoretic Approach. (Springer, New York, 1998).

30. Hastie, T., Tibshirani, R. & Friedman, J. The Elements of Statistical Learning. (Springer, 2016).

31. Benjamini, Y. & Hochberg, Y. Controlling the False Discovery Rate: A Practical and Powerful Approach to Multiple Testing. J. R. Stat. Soc. Ser. B 57, 289–300 (1995).

